# Phylogenetic and non-phylogenetic patterns in the richness, frequency and composition of links in a herbivore-parasitoid interaction network

**DOI:** 10.1101/2024.06.30.601383

**Authors:** Frazer H. Sinclair, Chang-Ti Tang, Richard A. Bailey, György L. Csóka, George Melika, James A. Nicholls, José-Luis Nieves-Aldrey, Alex Reiss, Miles Zhang, Albert Phillimore, Karsten Schönrogge, Graham N Stone

**Affiliations:** Institute of Ecology and Evolution, University of Edinburgh, Charlotte Auerbach Road, Edinburgh, EH9 3FL, Scotland, United Kingdom; Department of Ecology and Vertebrate Zoology, University of Lodz, Faculty of Biology and Environmental Protection, 12/16 Banacha St., 90-237 Łódź, Poland; National Agricultural Research and Innovation Centre, Forest Research Institute, Hegyalja Str. 18, Mátrafüred 3232, Hungary; Plant Health Diagnostic National Reference Laboratory, National Food Chain Safety Office, Budaörsi u. 141-145, 1118 Budapest, Hungary; CSIRO, Australian National Insect Collection, Clunies Ross Street, Acton, ACT 2601, Australia; Museo Nacional de Ciencias Naturales (CSIC), Departamento de Biodiversidad y Biología Evolutiva, C/ José Gutiérrez Abascal 2, ES-28006 Madrid, Spain; UK Centre for Ecology & Hydrology, Maclean Building, Benson Lane, Wallingford, OX10 8BB, UK

## Abstract

Revealing processes that structure species interactions is central to understanding community assembly and dynamics. Species interact via their phenotypes, but identifying and quantifying the traits that structure species-specific interactions (links) can be challenging. Where these traits show phylogenetic signal, however, link properties may be predictable using models that incorporate phylogenies in place of trait data. We analysed variation in link richness, frequency, and species identity in a multi-site dataset of interactions between oak cynipid galls and parasitoid natural enemies, using a Bayesian mixed modelling framework allowing concurrent fitting of phylogenetic effects of both trophic levels. In both link incidence (presence/absence) and link frequency datasets, we identified strong signatures of cophylogeny (related parasitoids attack related host galls) alongside patterns independent of either phylogeny. Our results are robust to simulations of substantially reduced sample completeness, and are consistent with the structuring of trophic interactions by a combination of phylogenetically conserved and convergently evolving traits in both trophic levels. We discuss our results in light of phenotypic traits thought to structure gall-parasitoid interactions and consider wider applications of this approach, including inference of underlying community assembly processes and prediction of economically important trophic interactions.

## Introduction

Biological communities comprise sets of species that interact via processes along a continuum from antagonism (e.g. predation, parasitism, competition) to mutualism (e.g. pollination, seed dispersal, parasite removal) (Bascompte et al. 2006; Cagnolo et al. 2011; Peralta 2016; Caves 2021). Revealing why some species interact but not others is central to understanding community structure, assembly and dynamics, and remains a fundamental goal of ecology (Cattin et al. 2004; Singer and Stireman 2005; Cavender-Bares et al. 2009; Yeakel et al. 2012; Bramon Mora et al. 2020). At a community level, interactions can be summarised as networks of pairwise species links (Bascompte et al. 2006; Cagnolo et al. 2011). Species vary in three attributes of their link distribution: richness (how many species each is linked to), identity (which species they are linked to), and frequency (how often a link is realised, a measure of interaction strength) (Memmott et al. 1994; Yeakel et al. 2012; Maia et al. 2019; Braga et al. 2020; Heimpel et al. 2021). Species interact via phenotypes, and many links are mediated by key biological traits (Ehrlich and Raven 1964; Thompson 2005). Well-studied examples include the roles of pollinator tongue length and flower depth in pollination mutualisms (Darwin 1862; Whittall and Hodges 2007; Anderson and Johnson 2008), and of plant chemical defences and herbivore countermeasures in herbivory (Ehrlich and Raven 1964; Janz 2011). Identifying such traits facilitates study of the impacts of natural selection and coevolution in community assembly (Thompson 2005; Rezende et al. 2007a; Pearse and Hipp 2009; Janz 2011; Fontaine and Thébault 2015; Endara et al. 2018), and allows modelling of network structures based on traits rather than species, which can improve predictive power (Truitt et al. 2019; Marjakangas et al. 2022; Pinilla-Gallego et al. 2022).

However, in many systems the traits mediating species links are difficult to identify, or may not be known for all members of an assemblage (Belshaw et al. 2003; Gripenberg et al. 2019). Alternatively, if phylogeny is a valid proxy for functional trait variation then species’ link properties may be predictable from phylogenetic relationships in each trophic level (Ives and Godfray 2006; Ives and Helmus 2011; Peralta 2016; Poisot and Stouffer 2018; Gripenberg et al. 2019; Gallinat and Pearse 2021; Perez-Lamarque et al. 2022; Benadi et al. 2022). Where this assumption holds, more closely related predators will be more similar in traits that structure their links with herbivores, and *vice versa* (Ives and Godfray 2006; Losos 2008; Poulin et al. 2011; Stouffer et al. 2012; Naisbit et al. 2012; Ives 2022) (our study considers trophic links, but the same rationale applies to other link types). Where this is true, species links may be predictable by a statistical model that uses a phylogeny for each trophic level as a proxy for functional trait variation (Ives and Helmus 2011; Ives 2022; Benadi et al. 2022). Statistical frameworks incorporating phylogenies can therefore have applications in predicting link properties for unsampled species, such as the natural enemies of an invading plant or herbivore, or the potential non-target victims of an imported biocontrol agent (Ives and Godfray 2006; Pearse and Altermatt 2013, 2015; Davies 2021; Heimpel et al. 2021), and also in predicting (co)extinction risk (Rezende et al. 2007b). The extent to which link properties show phylogenetic or other patterning can also aid inference of diversification mechanisms (Janz 2011). For example, in antagonistic bipartite networks (plant-herbivore, herbivore-enemy), a strong co-phylogenetic effect - in which related species in one trophic level are primarily linked to related species in another trophic level (Fig 1.D) - is characteristic of escape-and-radiate and host tracking diversification models in which mediating traits are phylogenetically conserved (Janz 2011; Endara et al. 2017, 2018). In contrast, demonstration that links between species are highly non-random but not associated with phylogenetic relationships in one or both trophic levels suggests convergent evolution of mediating traits and may be associated with a greater frequency of host-switching (Janz 2011; Endara et al. 2017, 2018). A major attraction of phylogeny-based inference of species link distributions is that while estimation of relevant trait values often requires system-specific measurements from multiple living examples of a single life stage (Albert et al. 2011; Wong and Carmona 2021), location of a species in a molecular phylogeny only requires DNA from a single example of any life stage. DNA sequencing and phylogeny reconstruction protocols are increasingly straightforward even in non-model taxa (Wachi et al. 2018).

**Figure 1.**
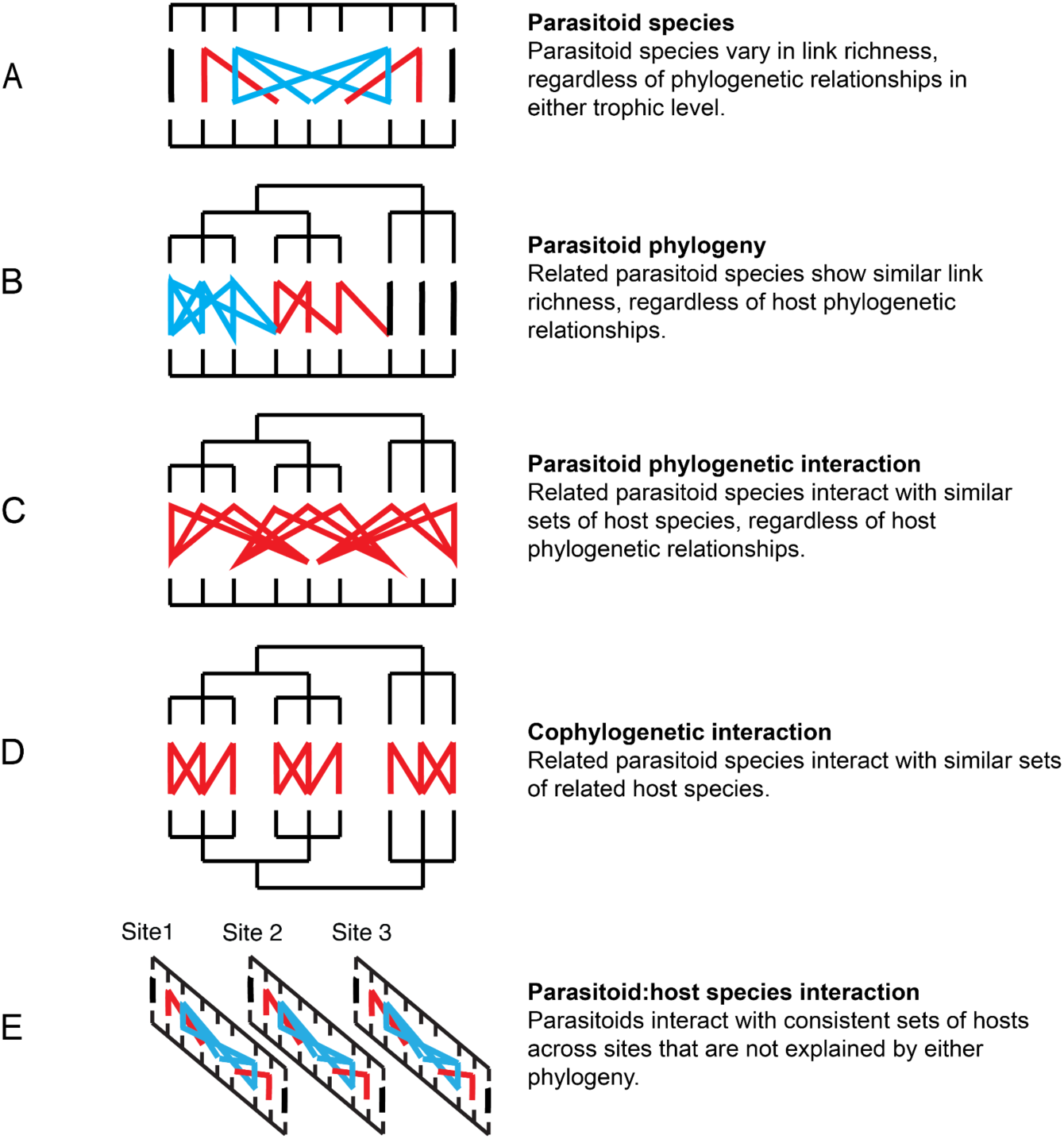
Five network patterns resolved as separate effects in a 2PhyGLMM for incidence data in MCMCglmm (Hadfield et al. 2014). Panels A and B show patterns in link richness (i.e. species vary in the number of taxa they interact with); panels C, D and E show patterns in link identity (species (co)vary in which taxa they interact with). Each diagram shows links (coloured lines) between an upper parasitoid trophic level and a lower herbivore trophic level. Effects B, C and D that incorporate phylogenetic information for one or both trophic levels show a resolved phylogeny for the trophic level concerned. Effects A to C are illustrated in terms of effects for the upper (parasitoid) trophic level. We co-estimated analogous effects for the herbivore trophic level alongside the illustrated parasitoid effects. Colours show variation in link richness from the parasitoid trophic level perspective; blue indicates parasitoids with 3 links (most generalist), red indicates 2 links, and black indicates parasitoids with a single link (most specialist). For clarity, potential links between some taxa in panels A and E have been omitted.

Despite these potential applications and the abundance of trophic link data (Poelen et al. 2014), the nature and strength of phylogenetic and non-phylogenetic patterning in link distributions remains poorly understood in most biological systems. Most methods and empirical analyses have focussed on whether close phylogenetic relatives in one trophic level commonly associate with - or are similarly abundant on - close phylogenetic relatives in the other trophic level (co- phylogenetic effects; fig. 1D), without incorporating other phylogenetic and non-phylogenetic effects (fig.1) (Legendre et al. 2002; Hommola et al. 2009; Leppänen et al. 2013; Eklöf and Stouffer 2016; Poisot and Stouffer 2018; Russo et al. 2018; Braga et al. 2020; Cruz-Laufer et al. 2022). Because accurate estimation of any single effect is conditional on which others are co-estimated (Hadfield et al. 2014; Perez-Lamarque et al. 2022), disentangling multiple effects requires a modelling framework that estimates multiple link richness and identity effects simultaneously, while controlling for potential confounding effects (Rafferty and Ives 2013; Hadfield et al. 2014; Gallinat and Pearse 2021). Several linear mixed modelling (LMM) approaches and their generalised extensions (GLMM) - here collectively termed two phylogeny mixed models (2PhyGLMM) - have been developed that allow such co-estimation, and have been applied to links recorded in terms of incidence (a binary response, whether two species interact or not; Hadfield et al. 2014; Endara et al. 2018) and counts (how frequently two species interact; Rafferty and Ives 2013; Hadfield et al. 2014). Advantages of model-based approaches include ability to incorporate alternative error structures associated with different data types, and additional variables such as spatial structure in the data or species traits (Rafferty and Ives 2013; Hadfield et al. 2014). To date, however, very few interaction networks have been analysed using two phylogeny mixed models (for examples, see Rafferty and Ives 2013; Hadfield et al. 2014; Endara et al. 2018; Galen et al. 2019; Lajoie and Kembel 2021). It is also remains unclear how sample sizes (i.e. number of trophic link records for each species at each trophic level) and data completeness impact on the ability to detect a signal (Bersier et al. 2002; Olesen et al. 2011; Rivera-Hutinel et al. 2012; Hadfield et al. 2014; De Aguiar et al. 2019)

Here we model link properties in an antagonistic bipartite network comprising herbivorous oak gallwasps and their parasitoid natural enemies. Modelling of both phylogenetic and non-phylogenetic effects requires bipartite interaction data across multiple sites, and a phylogeny for each trophic level (Hadfield et al. 2014). Our analysis uses 27,445 records of 54 parasitoid species reared from 38,638 galls of 60 cynipid species, collected from six sites in Hungary. This system is representative of a portion of the tritrophic insect-plant communities that comprise more than 50% of terrestrial biodiversity (Smith et al. 2008; Novotny et al. 2010), and has several properties that make it well suited for analysis of patterns in network structure. First, the parasitoids that attack oak cynipid galls are almost all specialists of this system (Askew et al. 2013), allowing it to be considered in ecological isolation (Bailey et al. 2009). Second, both trophic levels show wide variation in link properties, including richness (also termed the degree, specialisation or generality of a parasitoid species and the vulnerability of a host; Schoener 1989; Bersier, Banasek-Richter; and Cattin 2002), identity (also termed host range or host repertoire; Braga et al. 2020; Heimpel, Abram, and Brodeur 2021) and frequency (Askew et al. 2013). Both the first and second properties are shared with parasitoid communities centred on other herbivore guilds (Askew 1980; Stireman and Singer 2003; Leppänen et al. 2013; Santos et al. 2022). Third, there are good reasons to predict both phylogenetic and non-phylogenetic patterns in link identity in this system. Previous research on oak gall communities has identified traits that structure trophic links by influencing the ability of parasitoids to exploit specific host gall types (Bailey et al. 2009). These include parasitoid ovipositor toughness and length, which determine the depth to which a female parasitoid can locate hosts in target galls (Askew 1965; Quicke et al. 1998; Egan and Ott 2007), and a suite of gallwasp traits that, for parasitoids, influence their availability (phenology, host plant association, gall location on the host plant), accessibility (gall structural defences), and resource quality (host size) (Bailey et al. 2009). For both trophic levels, some link-structuring traits are phylogenetically conserved (e.g. gallwasp-oak associations, some gall defences, ovipositor length in some parasitoid lineages) while others show extensive convergent evolution between more distantly related galls (e.g. gall location on the oak, some gall defences, ovipositor length across parasitoid lineages) (Quicke and Belshaw 1999; Cook et al. 2002; Stone and Schönrogge 2003; Stone et al. 2009; Nicholls et al. 2018; Elias et al. 2018). Phylogenetic and non-phylogenetic patterns of links are thus possible in either or both interacting trophic levels.

Here we address the following questions: (1) How much of the variation in link richness, frequency, and identity is explained by phylogenetic and non-phylogenetic effects? (2) Do inferences differ for networks based on incidence data (binary presence-absence, unweighted networks) and frequency data (count-based, weighted networks)? We directly compare the variances explained by different phylogenetic and non-phylogenetic terms for incidence and frequency versions of the same datasets. (3) Which trophic associations contribute to observed patterns? (4) How robust is our inference? We use a subsampling scheme to explore the impact of variation in sampling completeness(Goldwasser and Roughgarden 1997; Jordano 2016) on inference and predictive power. We compare models that incorporate alternative approaches to incorporating sample size as a measure of sampling effort (Gotelli and Colwell 2011; Chao et al. 2020), to modelling pools of available interacting species (Hadfield et al. 2014), and incorporating spatial variation in community structure (Thompson and Townsend 2005; Brimacombe et al. 2023).

## Materials and Methods

### Gallwasp - Parasitoid Community Networks

We reared parasitoids from galls on four oak species (*Quercus cerris* L.*, Q. pubescens* Willd.*, Q. robur* L.*, Q. petraea* (Matt.) Liebl.), sampled at six sites in Hungary (for locations see fig. S1 in the supplemental PDF) between 2000–2003 using methods detailed in Bailey et al. (2009). The data include published records for five sites described in Bailey et al. (2009), with additional data for further gallwasp species from these sites and for a sixth site (Köszeg) sampled in 2000. Most oak gallwasps have lifecycles involving alternating spring (sexual) and summer/autumn (asexual) generations, each of which develops in a gall with a specific structural phenotype on a specific organ (acorn, bud, catkin, leaf, root) on specific host oak taxa. Each site was visited repeatedly through each sampling year to allow sampling of each generation of available cynipid species. All trees were sampled for the same fixed period of time, such that the gall frequencies obtained are a measure of relative abundance. All galls were identified to generation and species, reared individually, and monitored for emerging insects for 2.5 years. Images of gall phenotypes are available at https://doi.org/10.1371/journal.pbio.1000179.s002 and in Roskam (2019). The specific host(s) of the parasitoids within each galls were not determined - they could have attacked the gall inducer, cynipid inquilines or each other; our data represent association networks (links between parasitoids and host galls) rather than a trophic network (links between parasitoids and the cynipid gall inducer). All parasitoids were identified to species by expert taxonomists using morphological keys (Bailey et al. 2009). Full lists of sampled gallwasp and parasitoid taxa and an additional comment on taxonomic resolution are provided in the supplemental PDF, tables S1 and S2

The two generations of the same gall wasp species have very different gall phenologies and morphologies, sometimes develop on different plant organs on different host plant taxa, and support very different parasitoid assemblages (Bailey et al. 2009; Askew et al. 2013; Sinclair et al. 2015). As in Bailey et al. (2009), we therefore analysed data for sexual and asexual generations separately. Gall-parasitoid interaction matrices are shown for each generation in fig. 2. Summaries of sample sizes, link richness and link frequency for all gall generations and parasitoid taxa are provided in tables S1 and S2 in the Supplemental pdf, and the link datasets are available from the Edinburgh DataShare repository (https://doi.org/10.7488/ds/7760). The sexual generation dataset comprises 255 distinct link types identified from rearing 18210 adult parasitoids of 46 species from 11791 galls of 26 cynipid species, while the asexual generation dataset comprises 439 link types identified from rearing 9235 adult parasitoids of 43 species from 26847 galls of 52 cynipid species.

**Figure 2.**
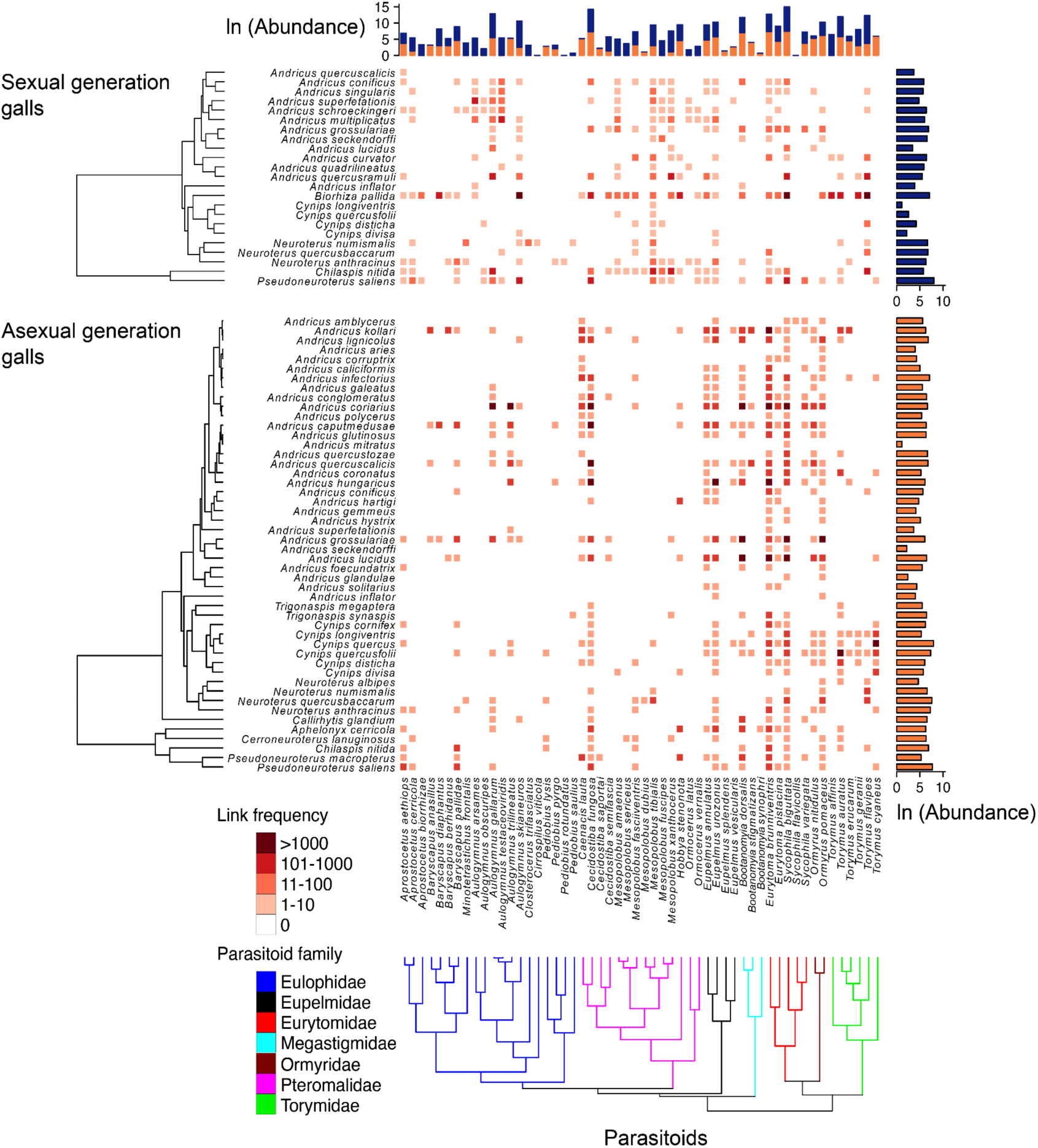
Heat maps showing bipartite link frequencies between parasitoids and sexual generation (top) and asexual generation (bottom) cynipid galls, pooled across sampling sites. Bar plots summarise sample sizes for host galls (right) and parasitoids (top) in the datasets for the sexual generation (blue) and asexual generation (orange). Phylogenies with node ages and branch support information are shown in Appendix 2.

For both generation datasets, we analysed link properties using incidence and frequency data. Incidence data incorporate no information on interaction strength (Berlow et al. 2004) and thus cannot discriminate between alternative scenarios in which a parasitoid species attacks two alternative hosts at relative rates of 99:1 and 50:50. In contrast, frequency data allow detection of cross-species variation in interaction strength. When designing sampling experiments, a valid question is whether additional inferences are possible from count data that justify the additional sampling effort required. The same phylogenetic and non-phylogenetic effects can be estimated in incidence and frequency models, though the interpretation of terms differs between them. For example, for incidence data the parasitoid phylogeny effect (fig. 1B) increases as related parasitoids attack an increasingly similar richness of host species, while for count data this occurs as related parasitoids inflict increasingly similar average attack rates on hosts. Although in this case the incidence-based interpretation is more straightforward, by not discriminating between weak and strong links (Berlow et al. 2004) a presence/absence approach may over-simplify patterns of enemy richness.

### Gallwasp and Parasitoid Phylogenies

We generated gallwasp and parasitoid phylogenies using partial sequences of one mitochondrial coding locus (cytochrome b for gallwasps and cytochrome c subunit 1 for parasitoids) and one nuclear non-coding locus (28S D2 for both trophic groups), totalling 999 and 1304 base pairs for gallwasps and parasitoids respectively. Full molecular and phylogenetic methods are provided in Appendix 2, and Genbank accession numbers of all sequences are provided in table S3 in the Supplemental PDF. To incorporate phylogenetic information into a GLMM, all root-tip distances are scaled to 1 and the inverse of the phylogenetic covariance matrix is generated for the gallwasp and parasitoid ultrametric trees (Hadfield 2010; Hadfield and Nakagawa 2010). To allow us to place observed phylogenetic patterns in a temporal context, we inferred node ages for both trophic levels (see Appendix 2). The phylogenies for each generation dataset used in MCMCglmm analyses are shown in simplified form in fig. 2, and are provided for all taxa in each trophic level and with full node age and branch support information in Appendix 2, figs. A1 and A2. Sequence alignments and treefiles for all three phylogenies have been deposited in the Edinburgh DataShare repository (https://doi.org/10.7488/ds/7760).

### Two Phylogeny Mixed Models

In 2014 Hadfield *et al*. developed and applied a GLMM approach to analysis of incidence data for links between fleas and their mammalian hosts. Independently, (Rafferty and Ives 2013) developed and applied an equivalent LMM to link frequency (count) data for insect visitors to flowering plants. Both approaches combine bipartite community interaction data with a phylogeny for each trophic level, and allow simultaneous estimation of variance components for link richness and link identity model terms, all of which are fitted as random effects. Frequentist forms (e.g. penalised quasi-likelihood) of two phylogeny mixed models can be fitted in software such as ASReml (Butler et al. 2017) or the phyr R package (Li et al. 2020).

However, while these are several thousand times faster than equivalent Bayesian approaches such as MCMCglmm, they behave poorly for Bernoulli data (as for incidence networks) where the grouping factors include phylogenetic or pedigree data, with likely downward bias in estimation of variance components (Gilmour et al. 1985; Hadfield et al. 2014). We therefore used the Bayesian MCMC framework incorporated in the package MCMCglmm (v2.32) (Hadfield 2010) in R (v4.1.0) (R Core Team 2021). R code for our analyses (including intraclass correlation calculations - see below - and figure plotting) is available . We summarise the terms fitted in our MCMCglmm models below, largely following (Hadfield et al. 2014), and give their biological interpretation in the context of host-parasitoid systems. Selected terms are illustrated diagrammatically for an incidence model in fig. 1 and all terms are described in full for both incidence and frequency models in Appendix 1. The patterns expected in a host-parasitoid link matrix when each term is substantial are illustrated in (Hadfield et al. 2014).

#### (a) Patterns in Link Richness

Patterns in link richness (species degree, for incidence data) and average link frequency (interaction strength, for count data) are captured by four random main effect (i.e. non-interaction) terms. Explained variance in richness/frequency across parasitoid and host species that lacks any phylogenetic pattern is allocated to a Parasitoid species effect (fig. 1A) and a Host species effect, respectively. The equivalent phylogenetically patterned effects are the Parasitoid phylogeny effect (fig. 1B) and the Host phylogeny effect, which in incidence models capture, respectively, the extent to which related sets of parasitoids attack a similar richness of hosts, and sets of related hosts are attacked by a similar richness of parasitoids. In frequency models, these effects capture among-species variation in average link frequency (average interaction strength, i.e., the extent to which parasitoids are rare to abundant across host species, and the extent to which hosts are rarely to heavily attacked across parasitoid species). Incorporation of datasets for multiple discrete samples (sites in our analysis) allows estimation of additional model terms. A Site main effect captures among-site variation in the proportion of realised links (equivalent to unweighted connectance) in an incidence model, and in average link frequency in a frequency model. The Site:parasitoid and Site:host interaction effects capture among-site variation in the richness of parasitoid and host species in incidence models, and in average link frequency in frequency models.

#### (b) Patterns in Link Identity

Patterns in link identity (i.e. the identity of the species forming links) are captured by four random effect interaction terms between hosts and parasitoids. The Parasitoid:host species interaction captures the extent to which specific sets of parasitoids attack (incidence model) - or have similar link frequency with (frequency model) - specific sets of hosts across samples, regardless of phylogenetic relationships in either trophic level (fig. 1E). This is equivalent to the species interaction of (Hadfield et al. 2014), renamed here to underline the involvement of both trophic levels, and can only be directly estimated in models that utilise information from patterns in species interactions across samples (here, across sites). Three additional terms capture the extent to which variation in link identity is predicted by one or both phylogenies. For the incidence model the Parasitoid phylogenetic interaction captures the extent to which related parasitoids attack similar sets of unrelated hosts (fig. 1C), the Host phylogenetic interaction captures the extent to which related hosts are attacked by similar sets of unrelated enemies, and the Cophylogenetic interaction captures the extent to which related taxa in one trophic level link with sets of related taxa in the other (fig. 1D). For the frequency model the same phylogenetic interactions are instead framed in terms of average link frequency (interaction strength). These three types of phylogenetic interactions are equivalent to the parasite evolutionary interaction, host evolutionary interaction and coevolutionary interaction of (Hadfield et al. 2014). We prefer our terminology because we are referring only to patterns in data, while use of the term coevolutionary implies additional demonstration of reciprocal adaptation (Althoff et al. 2014; Poisot and Stouffer 2018), which the model does not estimate.

### Model Fitting

Our study focuses on four core analyses: of incidence and frequency data, in both sexual and asexual generation bipartite networks. Our modelling of the above terms followed (Hadfield et al. 2014), as detailed in their equation 3 for phylogenetic terms (i.e Parasitoid phylogeny effect, Host phylogeny effect, Parasitoid phylogenetic interaction, Host phylogenetic interaction and Cophylogenetic interaction), and their equation 5 for non-phylogenetic equivalents (i.e. Parasitoid species effect, Host species effect, and Parasitoid:host species interaction). Modelling of sample sites followed treatment of regions by (Hadfield et al. 2014), with Site included as a random effect (as exemplified in their equation 7). We included two additional random effect terms not mentioned in (Hadfield et al. 2014) - the Site:parasitoid interaction and Site:host interaction - to account for variation in the richness (incidence models) and mean link frequency (frequency models) of hosts and parasitoids among sites. For ease of reference, interpretations of all fitted terms for incidence and count-based models are summarised in Appendix 1.

We modelled the sets of species available to form links at each site in each of two ways. In option 1, we assumed that the species recorded at each site in each gall generation represented all species available to interact locally and that the absence of a species is uninformative about species traits and biological processes governing community assembly (for instance absence may be due to biogeographic processes). Host and parasitoid links that did not occur because the two species were not present in the same site and generation were identified as structural zeros and removed from the dataset (J. D. Hadfield et al. 2014)). In option 2, we assumed that the full set of parasitoid species recorded in a given generation across all six sites was available to interact at each site, such that the absence of an interaction is informative about the focal processes underpinning community assembly. In this version of the dataset all links absent from a single site and generation were given a value of zero (i.e. structural zeros were included). Our rationale is that the ecological reality is somewhere between these two models, such that by fitting them both we can assess the sensitivity of our inference to which one is true. In practice, the two options produced very similar results for three of the four data sets (Supplemental PDF, tables S4, S5), though option 2 models took much longer to run. We therefore report results using option 1 and highlight differences between options where these arise. For each dataset, we also fitted a model in which the six site datasets were pooled into a single regional dataset. This requires option 2, and necessarily prevents fitting of model terms dependent on between-site variation. Comparison of these models with their full option 2 equivalents allows assessment of the sensitivity of other model terms to whether site level variation in link properties is accounted for.

We used a binomial model with logit link function for incidence data, and a Poisson model with log-link function for frequency (count) data. For the incidence models we followed (Hadfield et al. 2014) and fitted the logarithms of site-specific sample sizes for both hosts and parasitoids as fixed effects to account for the expectation that greater sampling of a species would lead to more of its links being observed. These terms were not included for the count-based models, which focus on the relative frequencies of different links rather than their presence-absence (Hadfield et al. 2014).

Parameter expanded priors were used for all random effect variances in both incidence and frequency-based models, with numerator and denominator degrees of freedom set to 1 and a scale parameter of 10^3^. For incidence based models the additive overdispersion term was fixed at 1, and for frequency-based models its prior followed an inverse Wishart distribution with nu = 0.002. Chains were run for 5 million iterations with a burn-in of 1 million and a thinning interval of 2000, resulting in 2000 sets of parameter estimates. We assessed the relative importance and statistical support for model terms using intraclass correlations (ICCs) on the latent scale (Nakagawa and Schielzeth 2010). We report the median and mode for ICC posterior distributions to indicate the relative contribution of model terms, based on 2000 posterior samples for each model, and interpret a term as significant where the lower bound of the 95% credible interval is removed from zero (defined here as > 0.01, as in (Hadfield et al. 2014)). MCMCglmm fits additive overdispersion models. For Poisson models the ICC denominator for each set of estimates included the sum of the random effect variances together with a residual that consists of an additive overdispersion term (i.e. the variance from an observation-level random effect) and a distribution specific variance, following (Nakagawa and Schielzeth 2010). For binomial models with a logit link the additive overdispersion term variance is not identifiable and was fixed at 1, with an added distribution-specific variance of π^2^/3 employed to estimate ICCs. Although these binomial models included fixed effects to control for sample sizes, the ICCs were ‘adjusted’ in that the fixed effect variances were not included in the ICC denominator (Nakagawa et al. 2017). For Poisson models with a log-link the additive overdispersion term was estimated from the data and the added distribution specific variance for ICC calculation was ln(1/exp(intercept))+1. To identify which host and parasitoid lineages contributed to each term’s ICC we calculated the posterior modes of the predicted MCMCglmm solutions either for individual species (as appropriate for the Host species, Parasitoid species, Host phylogeny and Parasitoid phylogeny terms), or pairwise links (as appropriate for the Parasitoid:host species interaction, Host phylogenetic interaction, Parasitoid phylogenetic interaction, and Cophylogenetic interaction terms). These solutions, also sometimes called true effect sizes, are similar to the best linear unbiased predictors (BLUPs) from frequentist mixed models (Sorensen 2009; e Silva et al. 2013).

We also used ICCs to summarise the overall explanatory power of our models (ICC_GLMM_), the magnitude of the three phylogenetic effects in link identity combined (ICC_PHY_), and the relative magnitude of phylogenetic versus non-phylogenetic link identity effects (ICC_REL-PHY_). For ICC_GLMM_, following (Nakagawa et al. 2017) the numerator for each of the 2000 posterior samples was the sum of variance parameters for all the fitted random effect terms and the denominator was the numerator plus a residual. For ICC_PHY_ the numerator was the sum for each set of posterior samples of the relevant phylogenetic link identity model terms (i.e., Parasitoid phylogenetic interaction, Host phylogenetic interaction and Cophylogenetic interaction) and the denominator was the sum of all the fitted random effect terms plus a residual. For ICC_REL-PHY_ the numerator was the sum of the relevant phylogenetic link identity model terms and the denominator was the numerator plus the non-phylogenetic link identity term (i.e. the Parasitoid:host species interaction).

### Sensitivity of Inference to Sample size

To assess the extent to which incorporation of sample size information influences the magnitude of model random effects, we compared the results of our full incidence models with alternatives without site-specific sample sizes as a covariate. This matters because in some systems, non-phylogenetic and/or phylogenetically-conserved variation in species abundance is thought to have causal impacts on associated link richness (Vazquez et al. 2005). It is thus unclear to what extent variation in gall sample sizes represents variation in sampling effort versus variation in a biologically relevant host trait. Where the latter is true, controlling for sampling effort using link frequency could reduce support for MCMCglmm random effects capturing patterns in link richness (Hadfield et al. 2014).

### Subsampling and Sample Completeness

All empirical trophic link datasets are likely to suffer from incomplete sampling, in that additional sampling could result in changed estimates of link incidence and/or frequency (Chao et al. 2020). Most common network metrics are sensitive to sampling completeness, though those incorporating interaction strength are usually less affected than those based on incidence (Nielsen and Bascompte 2007; Chacoff et al. 2012; Rivera-Hutinel et al. 2012; Vizentin-Bugoni et al. 2016; Falcão et al. 2016; Henriksen et al. 2019). A key question is then the extent to which results and inferences are sensitive to variation in sampling effort. We hypothesise that two phylogeny mixed models should show reduced ability to detect patterns in under- sampled networks (i.e. higher type II error) due to reduced host and parasitoid richness in incidence models, and that, as for network metrics, the inferences from count-based models should be less sensitive to reduced sampling than their incidence equivalents.

To quantify the impact of sample completeness on our inference, we generated datasets comprising 5%, 10%, 25%, 50%, and 75% subsets of the total number of sexual and asexual generation galls, based on a random draw process without replacement. These correspond to sample sizes of 590, 1180, 2948, 5895, and 8843 galls for the sexual generation datasets and 1342, 2685, 6712, 13424, and 20135 for the asexual generation datasets. To retain the methods and scope of the study but simulate a reduction in collecting effort, random draws treated each individual gall as a distinct sample without regard to species, site, or year of collection. At each subsampling level we generated 60 subsampling replicates that were used to fit 30 replicate incidence models and 30 replicate frequency models for both the sexual generation and asexual generation gall datasets, applying the same model structures used for the full models. As previously, the only zeros included in the resulting dataset were those for host-parasitoid species pairs that were both present in the same site and generation in the sub-sample. To assess the completeness of our sampling and to summarise the impacts of reduced sampling on the datasets, we calculated the Chao-2 and first order Jackknife (Jack-1) richness estimators (Gotelli and Colwell 2011; Chao et al. 2020) for host gall types, parasitoid species and pairwise interactions at each level of subsampling. Sampling completeness of subsampled datasets was summarised as the value of the metric obtained as a proportion of the estimated total richness for the full dataset. Both estimators give very similar results and for brevity we present results for the Jack-1 estimator.

## Results

Results for our four core analyses - incidence and frequency models for sexual and asexual generation gall communities - are summarised in table 1. Incidence and frequency model results for the same dataset were very similar, so we present them in parallel and highlight contrasts. All four models have substantial explanatory power on the latent scale, with median ICC_GLMM_ values (and 95% credible interval) ranging from 0.753 (0.671–0.860) for asexual generation frequency data to 0.829 (0.752–0.903) for sexual generation incidence data (table 1). In what follows, we identify the model term relevant to each inference in brackets, and in all cases inference is conditional on the other terms fitted in the model.

**Table 1.** Intraclass correlations (ICCs) for terms in the 4 core models (incidence and frequency- based models of sexual and asexual generation gall datasets). Values are presented as median/mode (and 95% credible interval) over 2000 MCMC point estimates. Significantly non-zero terms (i.e. where the lower bound of the 95% credible interval >0.01) are highlighted in **bold** for each model. ICC_GLMM_ is the ICC for all fitted model terms (i.e. excluding the residual), analogous to an R^2^ for the model. ICC_PHY_ is the combined ICC for phylogenetic interaction terms in link identity, and ICC_REL-PHY_ is the relative contribution of these terms to the ICC for all interaction terms in link identity.

| Term | Sexual generation galls |  | Asexual generation galls |  |
| --- | --- | --- | --- | --- |
|  | Incidence model | Frequency model | Incidence model | Frequency model |
| Site | .014 / .000<br>(.000-.096) | .027 / .016<br>(.000-.134) | .021 / .001<br>(.000-.126) | .037 / .023 (.002-.159) |
| Site:host interaction | .030 / .031<br>(.006-.069) | **.064 / .055**<br>**(.025-.122)** | **.033 / .029**<br>**(.012-.059)** | **.122 / .127 (.072-.175)** |
| Site:parasitoid interaction | .001 / .000<br>(.000-.006) | .002 / .000<br>(.000-.009) | .001 / .000<br>(.000-.005) | .003 / .000 (.000-.007) |
| Host species | .031 / .001<br>(.000-.142) | .024 / .001<br>(.000-.157) | .030 / .030<br>(.000-.075) | .061 / .069 (.000-.122) |
| Parasitoid species | .007 / .000<br>(.000-.049) | .040 / .039<br>(.000-.099) | .008 / .000<br>(.000-.052) | .032 / .001 (.000-.093) |
| Host phylogeny | .179 / .002<br>(.000-.471) | .314 / .340<br>(.000-.579) | .042 / .001<br>(.000-.202) | .061 / .002 (.000-.306) |
| Parasitoid phylogeny | .019 / .000<br>(.000-.106) | .015 / .001<br>(.000-.099) | .011 / .001<br>(.000-.086) | .013 / .000 (.000-.104) |
| Parasitoid:host species interaction | **.207 / .224**<br>**(.095-.329)** | **.143 / .159**<br>**(.068-.230)** | **.092 / .080**<br>**(.025-.164)** | .035 / .040 (.009-.068) |
| Host phylogenetic interaction | .013 / .000<br>(.000-.077) | .009 / .000<br>(.000-.070) | .024 / .001<br>(.000-.175) | .099 / .001 (.000-.231) |
| Parasitoid phylogenetic interaction | .133 / .123<br>(.000-.268) | .047 / .000<br>(.000-.110) | .040 / .000<br>(.000-.106) | .022 / .014 (.000-.049) |
| Cophylogenetic interaction | .089 / .002<br>(.000-.281) | .038 / .001<br>(.000-.192) | **.367 / .392**<br>**(.178-.537)** | **.185 / .142 (.026-.329)** |
| Residual | .171 / .161<br>(.097-.248) | .185 / .237<br>(.052-.299) | .244 / .258<br>(.172-.319) | .247 / .268<br>(.140-.329) |
| ICC denominator | 25.157 / 22.228<br>(16.192-40.403) | 30.150 / 27.075<br>(18.451-49.273) | 17.549 / 16.641<br>(12.996-24.187) | 26.786 / 26.672<br>(19.83-38.20) |
| ICC<sub>GLMM</sub> | **.829 / .839 (.752-.903)** | **.815 / .763**<br>**(.701-.948)** | **.756 / .742**<br>**(.681-.828)** | **.753 / .732**<br>**(.671-.860)** |
| ICC<sub>PHY</sub> | **.263 / .291**<br>**(.098-.443)** | **.118 / .076**<br>**(.033-.261)** | **.457 / .450**<br>**(.295-.600)** | **.318 / .309**<br>**(.196-.437)** |
| ICC<sub>REL-PHY</sub> | **.561 / .581**<br>**(.311-.763)** | **.460 / .445**<br>**(.186-.699)** | **.832 / .845**<br>**(.690-.957)** | **.900 / .889**<br>**(.807-.987)** |

### Patterns in Link Richness and Frequency

Neither non-phylogenetic (Host species effect, Parasitoid species effect) or phylogenetic (Host phylogeny effect, Parasitoid phylogeny effect) model terms for link richness and frequency were significant. While intraclass correlations (ICCs) were substantial for the host phylogeny effect in the sexual generation dataset (median ICC=0.179 for incidence data, 0.314 for frequency data; table 1) the lower 2.5% credible interval in both cases abutted zero and was non-significant by our threshold criterion. There was little evidence of among-site variation in the proportion of realised links or average link frequency (Site effect), or in parasitoid richness or average abundance (Site:parasitoid interaction in all four core models). However, host galls showed significant idiosyncratic among-site variation in richness for asexual generation galls (Site:host interaction, incidence data) and in average abundance for both generations (Site:host interaction, frequency data).

### Patterns in Link Identity

Phylogenetic effects in link identity were substantial in all four core models, with median (and 95% credible interval) values of ICC_PHY_ for combined phylogenetic terms ranging from 0.118 (0.033–0.261) in the sexual generation frequency model to 0.457 (0.295–0.600) in the asexual generation incidence model (table 1). Point estimates for ICC_PHY_ (i.e. overall phylogenetic signal) were greater for asexual than for sexual generation models (p=0.051), and greater for incidence than for frequency models (though ICC values in these different models cannot be formally compared). The contribution of phylogenetic terms to the ICC for combined phylogenetic and non-phylogenetic link identity effects, ICC_REL-PHY_, was always high but greater in the asexual generation than in the sexual generation for both incidence models (0.832 (0.690-0.957) versus 0.561 (0.311–0.763), p=0.011) and frequency models (0.900 (0.807–0.987) versus 0.460 (0.186–0.699), p=0.0005) (table 1).

Cophylogenetic interaction effects were large and significant in the asexual generation, indicating that closely related parasitoids commonly associate with (incidence) or are similarly abundant on (frequency) closely related host galls. This contrasts markedly with the pattern in the sexual generation, in which the cophylogenetic interaction term was non-significant (point estimate < 0.01) in both incidence and frequency models. The trophic links that make a strong contribution to cophylogenetic effects in the asexual generation are identified in figs. 3C and E. Hotspots (red) and coldspots (blue) correspond to phylogenetically related sets of parasitoids whose presence (incidence models) or frequency (frequency models) is predicted to be high or low, respectively, on phylogenetically related sets of hosts. Examples in our asexual generation data of hotspots include sets of links between: (i) *Andricus* gallwasps and a clade of generalist *Sycophila*, *Eurytoma* and *Ormyrus* parasitoids; (ii) *Andricus* gallwasps and *Eupelmus annulatus* and *E. urozonus*, (iii) *Cynips* gallwasps and *Torymus* parasitoids, and (iv) *Pseudoneuroterus* gallwasps and *Aprostocetus* parasitoids. In examples (i) and (iii), the parasitoids diversified substantially before the clade of host galls they attack (Appendix 2, figs A1 and A2). The parasitoid clade containing *Sycophila*, *Eurytoma* and *Ormyrus* (median age of most recent common ancestor with 95% posterior credibility interval = 50.1 (35.0-59.7) million years, MY) is substantially older than the *Andricus* clade whose galls they attack (MRCA = 10.2 (8.3-12.7)MY). Similarly, the *Torymus* clade (33.4 (26.3-42)MY) is substantially older than the *Cynips* clade whose galls they attack (5.7 (4.1-7.6) MY). In examples (ii) and (iv), the divergence of interacting host and parasitoid lineages is broadly contemporary. Divergence date estimates for *Eupelmus annulatus* and *E. urozonus* (14.8 (0- 27) MY) encompass those for the *Andricus* clade they interact with (see above), and the same is true for *Aprostocetus* parasitoids (3.0(0-12.0)MY) attacking *Pseudoneuroterus* galls (7.1(4.1-11.0)). Cold spots (coloured blue) indicate links whose absence (incidence models) or low frequency (frequency models) shows a strong phylogenetic pattern. Examples (particularly visible for frequency data in fig. 3E) include low levels of interaction between *Andricus* gallwasps and each of (i) *Aprostocetus* parasitoids, (ii) *Pediobius lysis,* and (iii) a clade of pteromalid parasitoids in the genera *Cecidostiba* and *Mesopolobus*.

**Figure 3.**
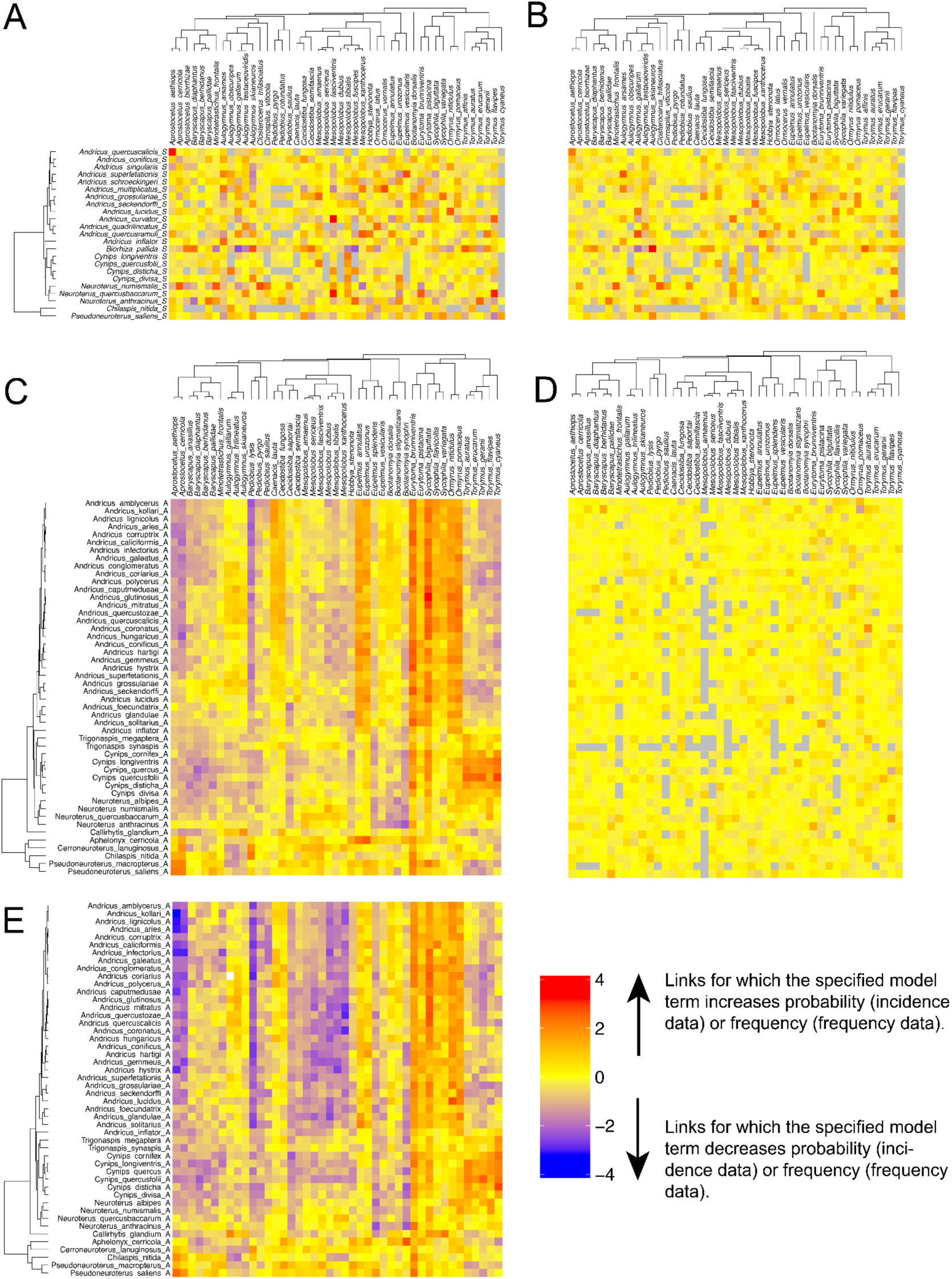
Heatmaps showing the extent to which specified model terms increase (red) or decrease (blue) link probability (incidence model) or frequency (frequency model). Grey-filled cells indicate host and parasitoid links whose solution was not estimated because the two species were not present in the same site and generation (i.e. structural zeros; see Methods). Cell values are posterior modes of the predicted MCMCglmm solutions (see Methods). In all plots, parasitoid taxa are arranged across the top, and cynipid gall generations down the side. A. Sexual generation incidence model parasitoid:host species interaction term. B. Sexual generation frequency model parasitoid:host species interaction term. C. Asexual generation incidence model cophylogenetic interaction term. D. Asexual generation incidence model parasitoid:host species interaction term. E. Asexual generation frequency model cophylogenetic interaction term. Gall taxon lists omitted from B and D are the same as those in panels to the left, and the parasitoid taxon list omitted from E is the same as for the panel above.

The two other phylogenetic effects in link identity (Parasitoid phylogenetic interaction and Host phylogenetic interaction) (table 1) were non-significant for all four core models. While ICCs were substantial for the Parasitoid phylogenetic interaction in the sexual generation incidence model (median ICC = 0.133) and for the Host phylogenetic interaction in the asexual generation frequency model (median ICC =0.099), in both cases the lower 2.5% credible interval abutted zero and thus neither was significant by our threshold criterion (table 1). The asexual generation frequency model including structural zeros for unsampled taxa (option b, see Methods) provided stronger support for a Host phylogenetic interaction (Supplemental PDF, table S5), indicating (in addition to a significant Cophylogenetic interaction) a further tendency for related hosts to be attacked by similar sets of unrelated parasitoids. The lack of individually significant phylogenetic terms in the sexual generation models despite high and significantly non-zero ICC_PHY_ values suggests that while there is substantial phylogenetic signal, it is not possible with our data to allocate it consistently to specific model terms.

Both generations showed additional patterning in link identity that was not associated with phylogenetic relationships in either trophic level (Parasitoid:host species interaction). This non-phylogenetic effect was supported in both incidence and frequency models and had higher ICC values for the sexual than for the asexual generation (table 1; the effect was marginally non-significant for the asexual generation frequency model by our threshold criterion). As we would expect for a non-phylogenetic effect, the strongly contributing links (dark red or blue in figs. 3A, B and D) are not phylogenetically clustered. We find many more strongly contributing links in models for the sexual generation (figs. 3A, B) than for the asexual generation (fig. 3D). The taxa involved in the strongly contributing links in the sexual generation are taxonomically diverse in both trophic levels, and again are very similar in incidence and frequency models (figs. 3A, B).

### Sensitivity of Inference to Sampling Effort and Pooling of data across sites

Removal of sampling effort as a covariate in incidence models did not change the most strongly supported model terms in either generation dataset. Models without sampling effort had generally higher ICCs for link richness model terms (for example, in the sexual generation the median ICC and 95% credible interval for the Host phylogeny effect increased from 0.179 (0.00-0.47) to 0.316 (0.00-0.59)), though these terms remained non-significant (see “Supplementary Results” in the Supplemental PDF; tables S4, S5).

For all four data sets, models in which data were pooled across sites showed elevated ICCs for most phylogenetic model terms relative to the equivalent full model (option 2; tables S4, S5), but no change in which of these were significant. Reallocation of site-associated variance to other terms in the pooled models was primarily apparent in elevated link richness terms. In the sexual generation frequency model the ICC for the Parasitoid species effect increased from a mean/mode (and 95% credible interval) of 0.054 / 0.047 (0.008-0.116, non-significant) to 0.112 / 0.084 (0.026-0.227, significant). In the asexual generation frequency model the ICC for the Host species effect increased from 0.047 / 0.034 (0.000-0.098, non-significant) to 0.141 / 0.132 (0.077-0.227, significant)

### Sensitivity of Model Results to Sampling Intensity

Our full datasets incorporated near complete sampling of the regional species pool for host galls (observed sexual and asexual generation richnesses are 100% and 98% respectively of the Jack-1 predicted values), but were less complete for parasitoid richness (86%) and interaction richness (75%) (See Supplementary Results in the Supplemental pdf; fig. S2). We used analyses of *in silico* subsampled datasets to assess the possible impact of incomplete sampling on our inference. Subsampling to 75%, 50%, 25%, 10%, and 5% of the complete datasets had little effect on host gall richness, which even for the 5% subsamples exceeded 80% and 88% of Jack-1 estimates for the complete sexual and asexual generation datasets, respectively. The effects of subsampling were more pronounced for parasitoid richness and link richness, which in the 5% subsamples (590/1342 galls for the sexual/asexual generation datasets) fell to around 50% and 20% of Jack-1 estimates for the complete data, respectively (fig. S2).

For most model terms identified as significant in the complete dataset, increasing levels of data reduction were associated with increasing variance in modal ICC estimates across replicate subsamples (fig. 4b), lack of significance in a growing proportion of them (fig. 4a), and hence an increasing type 1 (false negative) error rate. While the impact of data reduction on ICC variance was generally stronger in incidence than frequency models (as predicted), for most terms the consequences for significance, and hence inference, were similar for both data types (fig. 4a).

**Figure 4.**
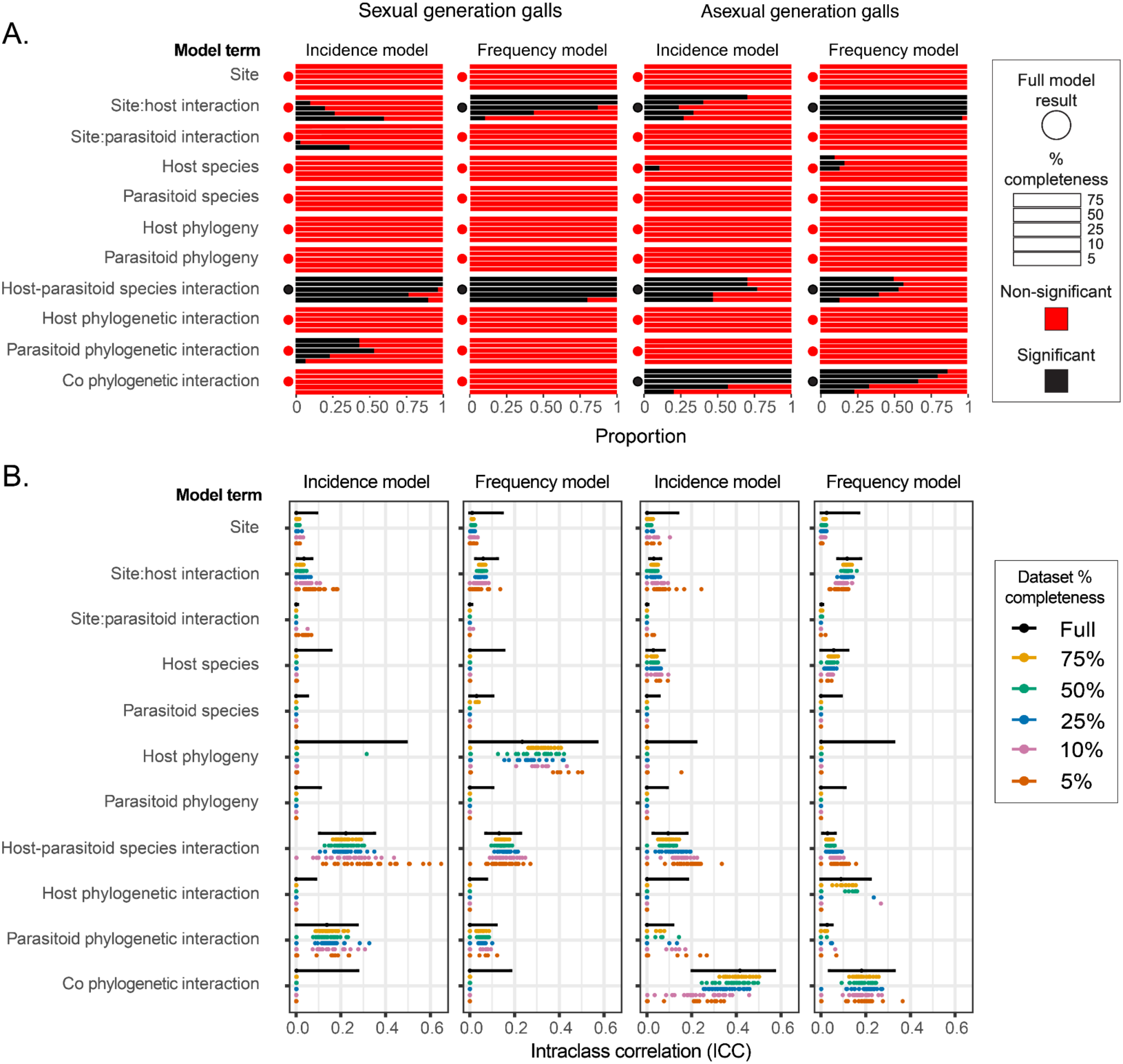
(a) Proportion of models for each generation and data type in which a given model term was significant (black bars: i.e. the lower bound of the 95% credible interval for the ICC >0.01) or nonsignificant (red bars) for different levels of data reduction. Filled circles to the left of each panel indicate the outcome for the model of the full dataset. Bars in each panel represent the proportion of significant and nonsignificant outcomes for each model term over 30 replicate data subsets for levels of data completeness from 75% (top), 50%, 25%, 10%, & 5% (bottom). (b) Intraclass correlations (ICCs) for models of parasitoid incidence and frequency on sexual and asexual generation galls. For models of full datasets, black dots and bars represent the posterior mode and 95% credible intervals for ICC estimates. For subsampled datasets, coloured dots represent the posterior modes for each of 30 replicate data subsets.

For both generations and data types, power to detect the dominant link identity terms in the full datasets remained high (i.e. supported in > 80% of subsampling replicates) with 50% data reduction (fig. 4a). For the sexual generation, power to detect the dominant Parasitoid:host species interaction in both datasets remained >75% even down to 5% subsampling. For the asexual generation, power to detect the dominant Co-phylogenetic interaction and the Host-parasitoid species interaction was more sensitive to data reduction in frequency rather than incidence models; power to detect the Co-phylogenetic interaction with incidence data remained 100% down to 25% sampling. Power to detect the Site:host interaction term (variation in host richness or average abundance between sites), significant in three of the four core full data models, declined in the sexual generation frequency and asexual generation incidence models, but remained close to 100% even at 5% subsampling in the asexual generation frequency model. In contrast, while the Site:host interaction term was not significant in the sexual generation incidence model for the full dataset, the number of model replicates in which it was significant increased with reduced sampling, reaching 50% for the 5% dataset.

## Discussion

We uncovered phylogenetic and non-phylogenetic patterns using a model-based approach to study trophic links between chalcid parasitoids and host oak cynipid galls. Our spatially replicated sampling allowed us to co-estimate effects on link richness (species degree), link frequency (interaction strength) and link identity that are independent of phylogeny in either trophic level, or associated with phylogeny in a specific single trophic level, or associated with both phylogenies. We analysed separate datasets for sexual (spring) and asexual (late summer, autumn) generation galls, which support substantially different parasitoid assemblages (see fig. 3 in (Bailey et al. 2009)). Our models had substantial explanatory power (median ICC_GLMM_ ≥ 0.753), and we obtained very similar sets of significant terms in models using incidence or frequency data, with or without sampling effort as a covariate, and using alternative approaches to modelling available pools of interacting species.

We found no significant phylogenetic or non-phylogenetic patterns in link richness (species degree) in incidence models or average link frequency (interaction strength) in frequency models. Thus we found no consistent tendency for related parasitoids to be similarly specialist or generalist, or to attack hosts at similar average rates, and no significant tendency for related host galls to be attacked by a similar richness of parasitoids, or to experience a similar mean parasitoid attack rate. Phylogenetic patterns in link richness and frequency are less well studied than patterns in link identity, but our findings are consistent with high evolutionary lability in host richness in *Eupelmus* (Al Khatib et al. 2016), one of the groups of chalcid parasitoids in our study. Our results also contrast with detection of phylogenetic signal in these properties in some mutualistic pollination and fruit dispersal networks (Rezende et al. 2007a; Tedersoo et al. 2013) and antagonist parasite-host networks (Poulin et al. 2011).

In contrast, we found strong phylogenetic and non-phylogenetic patterns in link identity. Asexual generation galls showed a dominant cophylogenetic signature, with related parasitoids tending to attack (and show similar attack rates on) sets of related host galls. The strength of phylogenetic signal in link identity, measured by our metric ICC_PHY_, was greater for the asexual generation than for the sexual generation, in which no single phylogenetic effect received consistent support. ICC_PHY_ was nevertheless also substantial and significantly non-zero for the sexual generation, suggesting that there was not enough information in this dataset to consistently partition substantial phylogenetic patterning amongst the phylogenetic terms. The cophylogenetic signal in the asexual generation parallels previous demonstration of similar signatures in host-parasitoid networks (Ives and Godfray 2006; Nyman et al. 2007; Leppänen et al. 2013; Wang et al. 2023), host-pathogen interactions (Clark and Clegg 2017), marine food webs (Eklöf and Stouffer 2016) and mutualistic plant-pollinator and plant-frugivore networks (Rezende et al. 2007b). The strongest pattern in sexual generation galls, however, was different: similar sets of unrelated parasitoids tend to attack (and show similar attack rates on) similar sets of unrelated host galls across sites. This same non-phylogenetic effect was also present, though less strongly, in asexual generation galls. To our knowledge this is the first study of herbivore-parasitoid communities to reveal such spatially consistent patterns in link identity, having controlled for phylogenetic and sample size effects (see (Endara et al. 2018) for a parallel application of MCMCglmm to interactions between *Inga* tree and insect herbivores across multiple sites in South America).

Previous studies have found greater phylogenetic conservatism in link identity for hosts/prey than for their parasites/predators (Ives and Godfray 2006; Naisbit et al. 2012; Peralta 2016; Cruz-Laufer et al. 2022). Were oak cynipid-parasitoid links to show similar asymmetry in phylogenetic signal between trophic levels, then having controlled for the Cophylogenetic interaction we would expect ICC support for the Host phylogenetic interaction. While we found no evidence of this in models including structural zeros (option 1), both the Cophylogenetic interaction and Host phylogenetic interactions were substantial and significant in Asexual generation models in which host-parasitoid interactions observed at any site were assumed possible at all sites (Option 2; Supplemental pdf, table S5). This is consistent with patterns in other systems. Peralta (2016) suggests that this patterns arises because traits determining enemy ability to exploit hosts or prey are expected to evolve faster than traits determining host/prey vulnerability, which should thus have a stronger phylogenetic signal in network structure (Rossberg et al. 2006; Fontaine and Thébault 2015). We might expect stronger host than parasitoid phylogenetic effects in the oak gallwasp system because while the hosts are a single radiating lineage, the parasitoids attacking each gall comprise representatives of several chalcidoid families that diverged before the origin of Cynipini (see below). Thus even if parasitoid lineages have tracked host diversification after their initial recruitment, there is a deep non-phylogenetic signal for this trophic level in this system.

### Host-Parasitoid Community Assembly

The patterns revealed by our models are generated by patterns of cross-species variation in traits that shape species interactions. These include those that predict species overlap in space and time (such as habitat, host plant, phenology, population size and geographic distribution) (Plantard and Hochberg 1998; Lindenfors et al. 2007; Slove and Janz 2011; Slatyer et al. 2013; Nicholls et al. 2018; Warren et al. 2022), and those that influence the outcome of encounters between species. For parasitoids, successful exploitation of a host gall requires behavioural and physiological traits (such as chemosensory systems) that allow detection of the gall and of hosts concealed within it, morphological traits (such as ovipositor length) that allow successful oviposition through gall tissues, and the ability to develop on the resources a host provides provides (Askew 1965, 1980; Quicke et al. 1998; Egan and Ott 2007; Bailey et al. 2009). For host galls, vulnerability/resistance depends on a suite of traits including gall morphological defences (Bailey et al. 2009; Egan et al. 2011), recruitment of ant bodyguards (Abe 1992; Inouye and Agrawal 2004; Warren et al. 2022), and chemical defences (Guiguet et al. 2023). Our contrasting results for sexual and asexual generation datasets could indicate structuring by different sets of traits, or contrasting evolutionary histories of the same (or overlapping) sets of traits. Between-generation differences must involve traits that are free to evolve separately in the two generations, but cannot be shaped by properties that both generations are constrained to share - such as habitat and geographic distribution. All of the gall traits thought to mediate cynipid interactions with parasitoids appear able to evolve independently in sexual and asexual gall generations, and show examples of both convergent evolution and phylogenetic conservatism within lineages (Stone and Cook 1998; Cook et al. 2002; Stone et al. 2009; Nicholls et al. 2017; Ward et al. 2022). While little is known about patterns of evolution in traits associated with host location behaviour in chalcidoid parasitoids, ovipositor length shows both phylogenetic conservatism and convergent evolution in this group (Al Khatib et al. 2016; Maletti et al. 2021). The best way to improve our models, and better understand underlying processes, is to incorporate information for the variables (including the candidate traits for both trophic levels above) that underlie the patterns captured by our random effects (Ives 2022). A strength of MCMCglmm and similar model-based approaches (Rafferty and Ives 2013; Ives 2022) is that incorporation of trait data is straightforward in principle, though the models may take a very long time to run. Persistently strong phylogenetic or non-phylogenetic model terms after inclusion of known candidate traits would be informative in implying either that the candidate traits have been correctly identified but incorrectly quantified, or that additional important structuring traits remain to be discovered. An example of this process, incorporating plant chemical defensive traits into MCMCglmm analysis of a tree-herbivore network, is provided by (Endara et al. 2018).

Our heat maps of predicted model solutions (fig. 3) show that the importance of significant model terms is not uniformly distributed across links in each network: instead, particular terms apply more or less for particular interactions. Notwithstanding this caveat, strong patterns in link identity in both gall generations that are uncorrelated with either the parasitoid or host gall phylogeny are compatible with ecological sorting of trophic interactions by convergently evolved traits. In contrast, the dominant cophylogenetic interaction in asexual generation galls confirms a major role for structuring by traits that are phylogenetically conserved in both trophic levels. Cophylogenetic patterns can arise through (a) coevolutionary codiversification, driven by arms race-type reciprocal adaptive change in gall defences and parasitoid countermeasures (Currie et al. 2003); (b) phylogenetic host tracking, in which parasitoids radiate across an existing diversity of hosts; (c) trait-based sorting of existing parasitoid lineages over a later radiation of hosts (ecological sorting) (Janz 2011, Althoff et al. 2014). The chalcid parasitoid lineages in our study diversified over 125 million years ago (Cruaud et al. 2024), long before cynipid gallwasp hosts were available (Blaimer et al. 2020). Assembly of this community has thus involved independent shifts onto gallwasp hosts by multiple parasitoid lineages over tens of millions of years, implying an initial role for ecological sorting (the same is true for parasitoids attacking sawfly hosts (Leppänen et al. 2013)). Concordant ages for some gallwasp and parasitoid divergence events (e.g. for *Eupelmus* parasitoids attacking *Andricus* galls and for *Aprostocetus* parasitoids attacking *Pseudoneuroterus* galls) are more compatible with codiversification.

### Food web prediction in the absence of trait data

Our compound predictors ICC_PHY_ and ICC_REL-PHY_ revealed that, despite lack of consistent support for specific terms in some datasets, our models explained high percentages of the variation in link properties, with phylogenetic effects being universally important. The lack of significance of individual predictors is therefore likely a consequence of insufficient informativeness (lack of orthogonality) in the data for separating out all the many random effect predictors included in the model. Despite this, the strong ICC_PHY_ and ICC_REL-PHY_ effects indicate high power to predict either gallwasp or parasitoid species interactions based on phylogenetic position in the absence of trait data (Ives and Godfray 2006; Poisot and Stouffer 2018; Braga et al. 2020, 2021; Strydom et al. 2022). For our data, ICC_PHY_ and ICC_REL-PHY_ values also indicate greater power to predict link identity in models based on incidence data rather than frequency data. Given the substantial effort required to sample interaction data, the extent to which interactions can be accurately predicted by trait-free phylogenetic models is an important question (Strydom et al. 2021, 2022). In networks with strong phylogenetic signal, such an approach has high potential applied value in relatively low cost prediction of interactions in invasive species, introduced control agents, and also species of conservation concern. The strong non-phylogenetic effects we observed can also have predictive power, but only for the specific species and sites present in the tested dataset. An interesting avenue for future research is to survey the variation in ICC_PHY_ and ICC_REL-PHY_ values across different types of interaction network. Such a comparative approach should be helpful in identifying key drivers of phylogenetic signal having controlled for other effects.

### Critique of our data and approach

Inference from two phylogeny mixed models is potentially sensitive to multiple aspects of the data used to construct the network and phylogenies, and the sampling design. We therefore assessed potential impacts of sampling completeness (Goldwasser and Roughgarden 1997; Jordano 2016), modelling of sample size (Gotelli and Colwell 2011; Chao et al. 2020), available species pools (Hadfield et al. 2014) and spatial structure (Thompson and Townsend 2005; Brimacombe et al. 2023). Here we consider these in turn, together with potential impacts of phylogenetic uncertainty (Perez-Lamarque et al. 2022).

#### (a) Sampling completeness

Analyses of simulated networks show that the sampling effort required to adequately estimate network properties is strongly dependent on sampling design and underlying network topology (De Aguiar et al. 2019). The same considerations are likely to influence MCMCglmm model-based inference. Analyses of replicate subsampled datasets showed that power to identify the same sets of significant model terms was largely resilient to at least 50% data reduction (fig. 4a). Such stability of inference implies that the patterns we observe in the full datasets are unlikely to be artefacts of undersampling.

#### (b) Sample size

We incorporated sample size as a covariate in incidence models to capture the potential impact of the numbers of each species sampled on the detection of links involving them (Nyman et al. 2007; Bailey et al. 2009; Hadfield et al. 2014; Endara et al. 2018). However, with a standardised sampling scheme (as here), interspecific variation in sample size likely reflects underlying variation in population size, a biological trait that can covary with phylogeny or niche (Vazquez et al. 2005). Incorporation of sampling effort thus has the potential to reduce support for other model terms (Hadfield et al. 2014). We assessed potential for this effect by comparing models with and without incorporation of sample size. For both generation incidence datasets, exclusion did not change the dominant model term, though a significant Parasitoid:host species interaction was lost for the asexual generation (table S5).

#### (c) Phylogenetic Uncertainty

We expect ability to accurately resolve phylogenetic patterns in link properties to depend on the strength of those patterns, the topology of the true phylogeny, and the accuracy with which empirically derived phylogenies capture true relationships (Ives and Godfray 2006; Hadfield et al. 2014; Ives 2022; Perez-Lamarque et al. 2022). In MCMCglmm, phylogenetic covariance between a pair of taxa is modelled as the proportion of total tree height that is shared from the root of the phylogeny to their most recent common ancestor. The greater this proportion, the greater the expected covariance (Hadfield 2010; Hadfield and Nakagawa 2010). Errors that distort the relative phylogenetic distances between taxa distort the covariance matrix, and hence potentially influence the outcome of MCMCglmm models. Uncertainty over relationships basal to the most recent common ancestor of a pair of taxa does not alter the branch length they share, and so has no direct impact on MCMCglmm models. We have high confidence in our gallwasp phylogeny (Appendix 2, fig. A1), which has high congruence in topology with a previous analysis using larger samples of taxa and hundreds of genes (Blaimer et al. 2020). Resolving chalcid parasitoid relationships is much more challenging, due to a widely recognised signature of rapid radiation (short internal branch lengths) towards the root of the phylogeny (Munro et al. 2011; Cruaud et al. 2024). We therefore used information from recent phylogenomic analyses to inform construction of our parasitoid phylogeny (Appendix 2, fig. A2). Our phylogeny resolves the same monophyletic families as a recent genome-level analysis with high support (Cruaud et al. 2024). Because our phylogenetic uncertainty for parasitoids largely concerns support for short internal branches deep in the phylogeny, we suggest that impacts on estimates of variance covariance, and hence on our inference, will be small. One could quantify the impact of phylogenetic uncertainty by fitting MCMCglmm models to sets of alternative phylogenies for one or both trophic levels (Healy et al. 2014). However, the computational effort required to do this in MCMCglmm would be prohibitive - it would take over 25 core years on the machine used in this study to do only a hundred replicates of each of the core models.

#### (d) Available Species Pools

A potentially important issue in modelling of species interactions concerns the interpretation of links that are present at some sites but not others. Each unobserved link could be genuinely absent, or be present but undetected (Olesen et al. 2011; Terry and Lewis 2020). We fitted alternative models for these situations in MCMCglmm (our options 1 and 2, respectively). For our data the two options gave the same dominant effects - the main difference being the significant support for a Host phylogenetic interaction in addition to a significant Cophylogenetic interaction only for option 2 in the Asexual generation frequency model. In the Western Palaearctic, the oak gallwasp system is characterised both by short-term patchiness in distributions and frequencies of interaction of individual species (captured by our option 1), and by geographically wide distributions and records of species interaction (captured by option 2) (Washburn and Cornell 1981; Askew et al. 2013). In each study system the truth is likely to lie somewhere between these two models, and the extent to which their results are concordant provides an indication of how sensitive inference is to model choice.

#### (e) Spatial Structure

Incorporation of site level information is fundamental to estimating non-phylogenetic patterns in link properties in our full MCMCglmm models, because this effect is estimated through consistency of interactions across replicate samples. Incorporation of site level information was particularly important for our Sexual generation models - we would have detected no significant patterns in link identity without it. Incorporation of site level information also allows controlling for spatial variation in abundance in both trophic levels, with the possibility of scoring unsampled interactions as structural zeros rather than real data zeros (see (d), above). The effect this can have on inference suggests that spatial structure should be incorporated where possible.

## Acknowledgements

This research was funded by NERC grants NE/T000120/1 to GNS, AP and KS, NE/E014453/1 to GNS and JAN, and GR/12847 to GNS and KS. YMZ was supported by the European Union’s Horizon 2020 research and innovation programme under the Marie Skłodowska-Curie grant agreement no. 101024056. Alex Reiss is funded by a PhD studentship from the UKRI-funded EASTBIO doctoral training program. For the purpose of open access, the authors have applied a Creative Commons Attribution (CC BY) licence to any Author Accepted Manuscript version arising from this submission.

## Appendix 1. Model terms and the corresponding interpretations if substantiated in incidence or frequency-based MCMCglmm models

A link is here defined as a bipartite association between specific host gall and parasitoid taxa. Host here refers to the gall from which a parasitoid emerged, rather than its trophic host. In incidence models, link richness (how many taxa a focal taxon is linked to) is also termed the degree, specialisation or generality of a parasitoid species, and the vulnerability of a host. In frequency models the equivalent terms capture variation in the mean frequency with which a focal taxon interacts with others, and are measures of interaction strength. Link identity (which species interact) is also termed host range or host repertoire. For the Site random effect, which links are considered possible depends on whether unsampled links are modelled as totally absent (structural zeros, option 1), or present but unrecorded (data zeros, option 2) (see Methods). Abbreviations in brackets after a term name are as used by Hadfield et al. (2014). Non-phylogenetic random effect terms that are marked with **†** incorporate an identity matrix for variance covariance between levels.

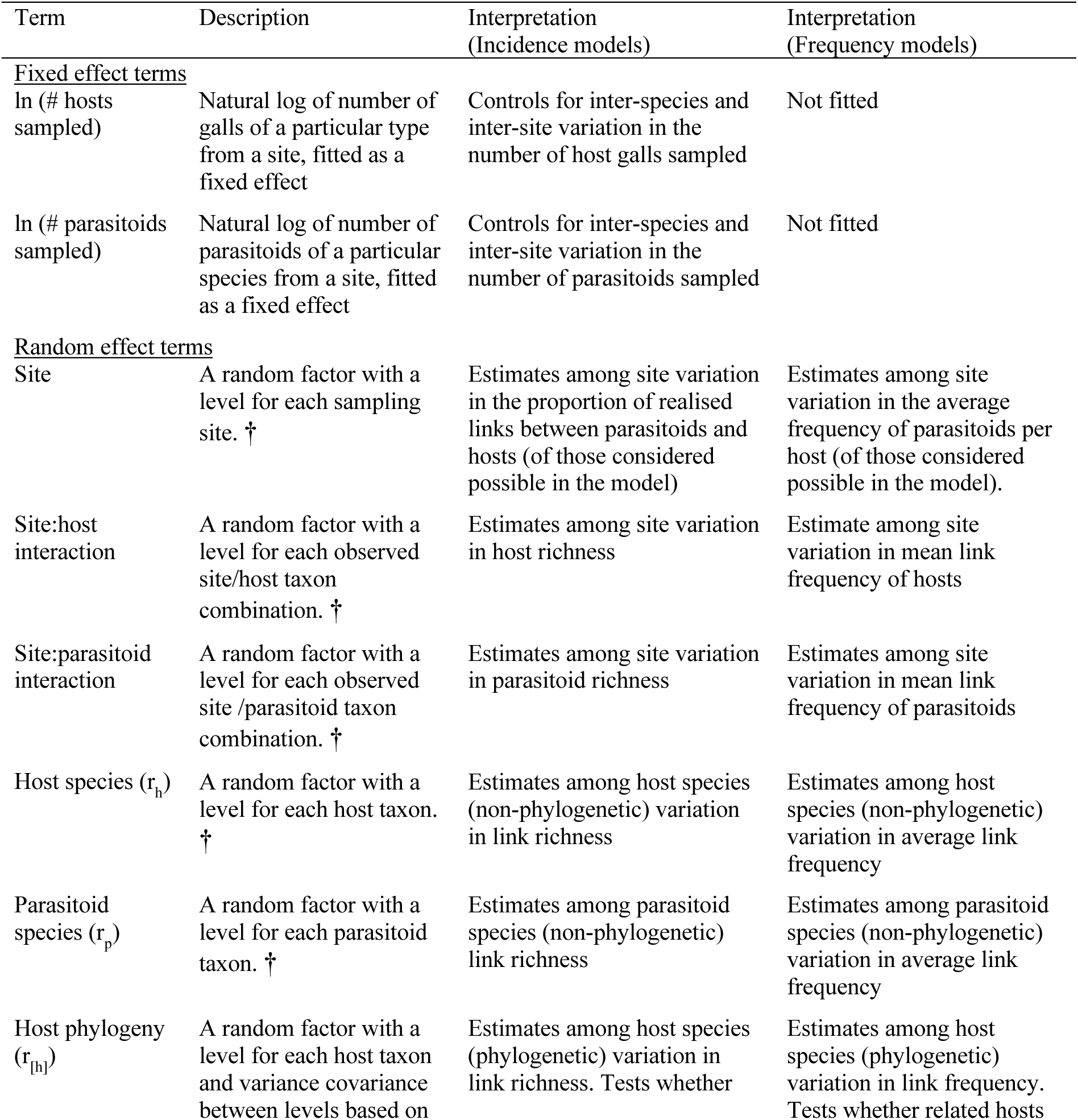
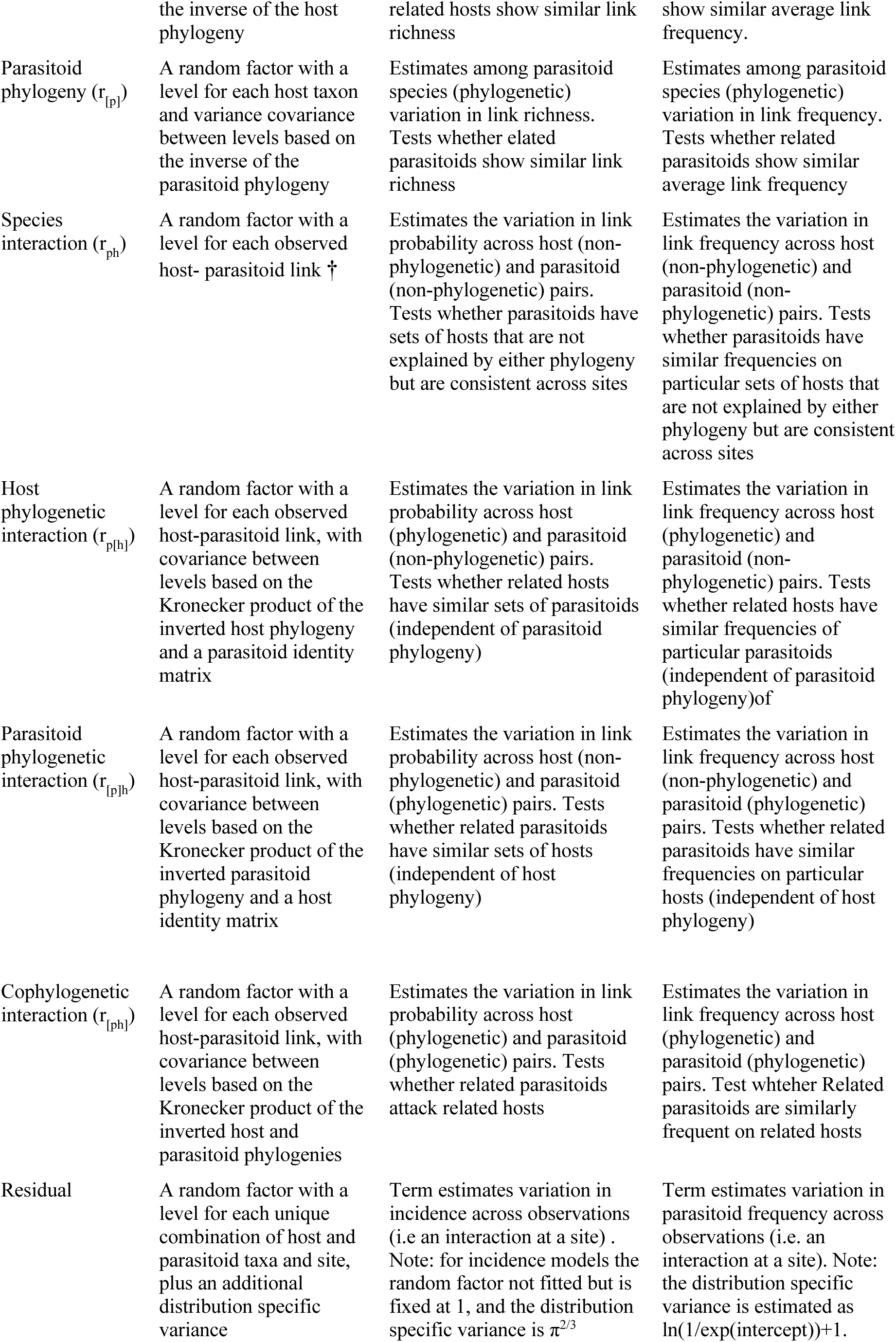

## Appendix 2. Phylogenies used in MCMCglmm analyses, and detailed molecular and phylogenetic methods

**Figure A1.**
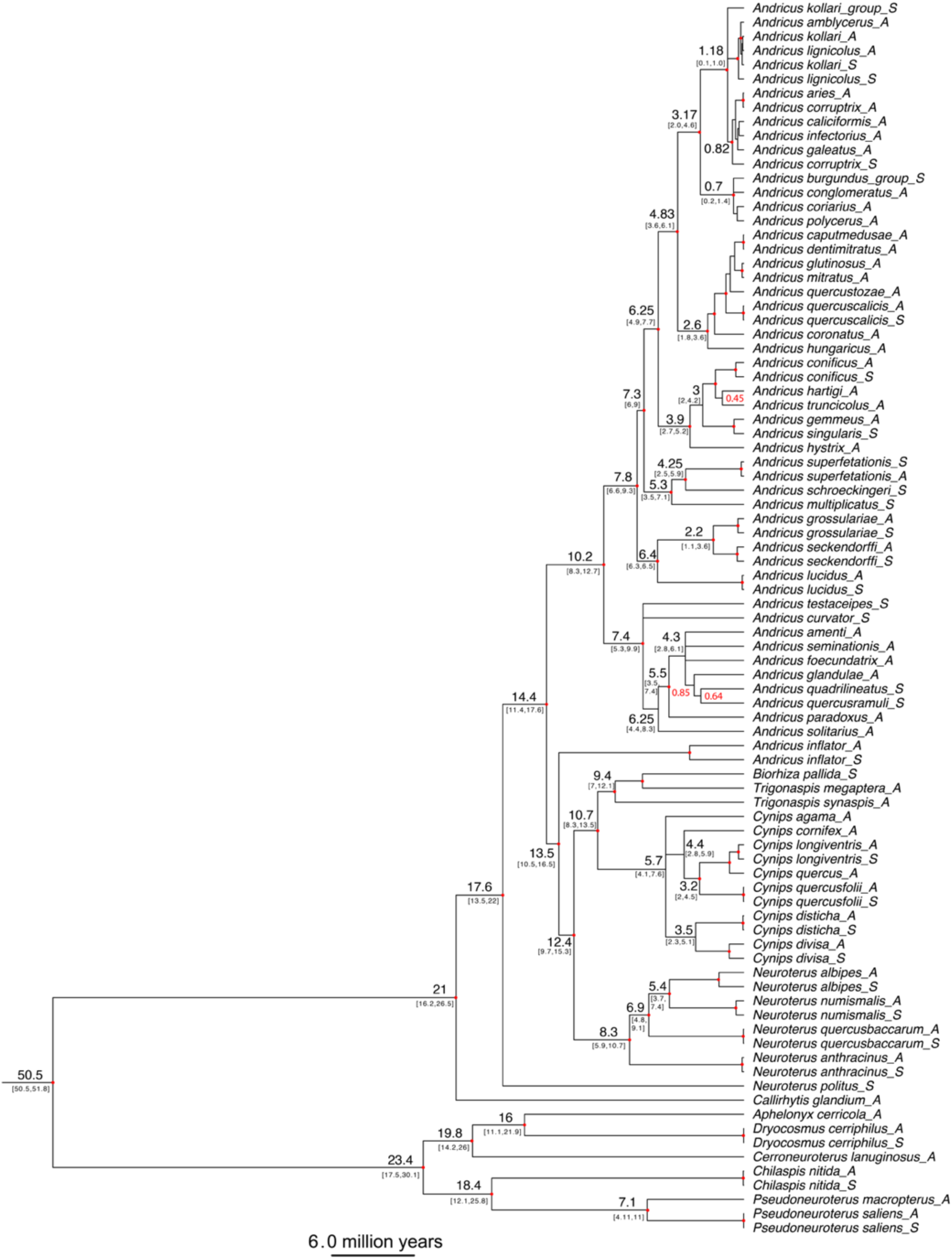
Maximum clade credibility tree for the cynipid gallwasps in this study used in MCMCglmm analyses. Numbers in black indicate node age in millions of years (with 95% posterior credibility interval). To preserve clarity of the figure, some node ages have been excluded. Posterior probability for all nodes is 1.0, with the exception of 3 nodes whose posterior probability is labelled in red, and for very recent divergences in the topmost clade containing *Andricus kollari*. All nodes marked with a red circle are shared with the best supported relaxed clock model, which also shows very similar branch length and node dates (data not shown).

**Figure A2.**
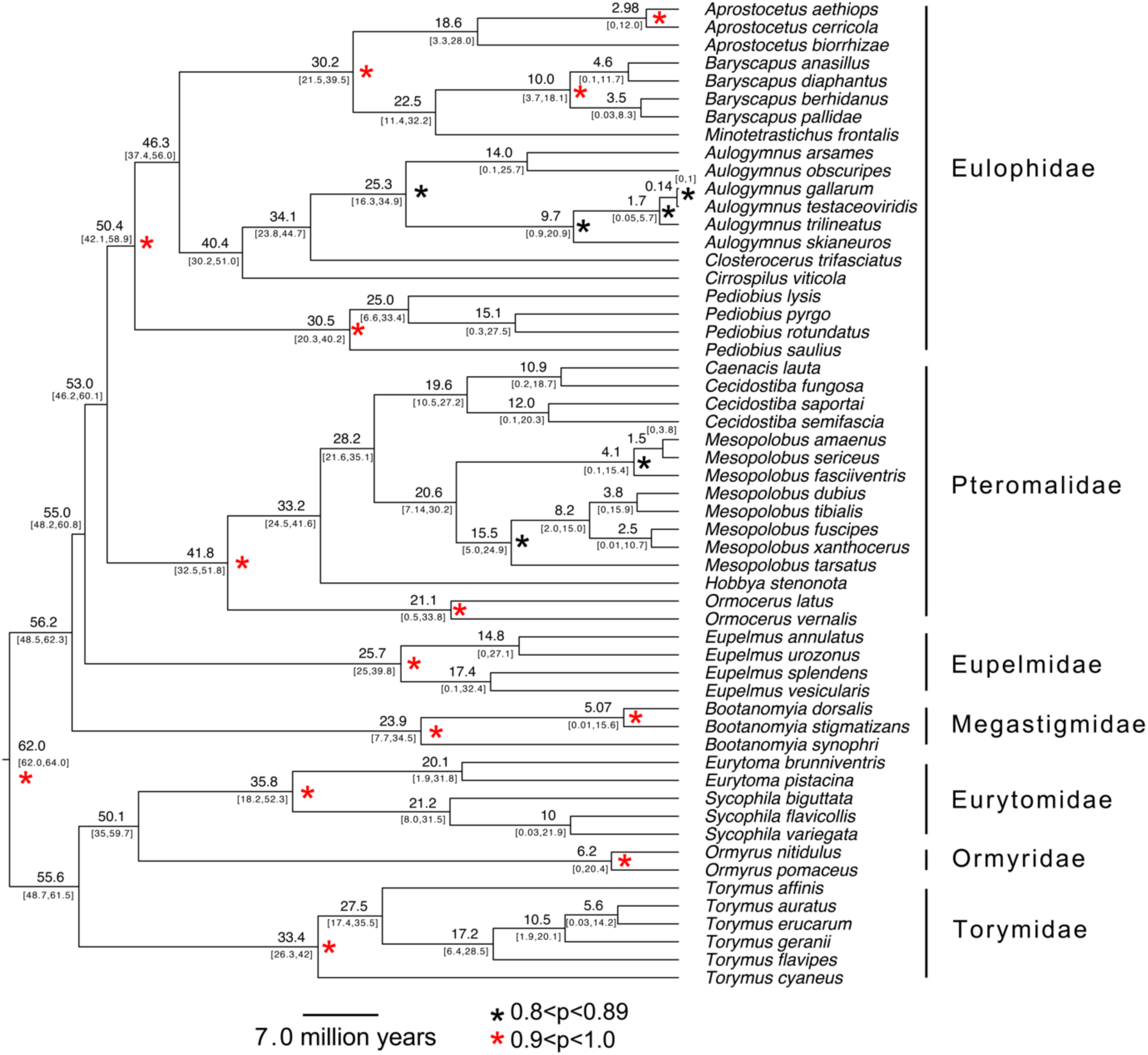
Maximum clade credibility tree for the chalcid parasitoids in this study used ibn MCMCglmm analyses. Numbers in black at selected nodes indicate node age in millions of years (with 95% posterior credibility interval). Our estimates for family common ancestors are close to the youngest inferred in a recent extensive analysis by Cruaud et al. (2024). Posterior probabilities of 0.90-1.0 are labelled with a red asterisk, and of 0.80-0.89 are labelled with a black asterisk. The family Eulophidae were constrained to be monophyletic in the analysis; monophyly was inferred for all other families with high posterior support.

### Detailed molecular and phylogenetic methods

#### Sequence generation and alignment

Phylogenetic relationships were reconstructed using fragments of one mitochondrial coding gene [698bp of cytochrome c oxidase subunit I (COI) for parasitoids, 433 bp of cytochrome b (cytb) for gallwasps] and a 560–580 bp long fragment of the non-coding 28S D2 region (both trophic levels).

Parasitoid COI sequences were amplified using primers COI_pf1: 5’ AGG RGY YCC WGA TAT AGC WTT YCC 3’ (designed by James Nicholls) and COI_2413d: 5’ GCT ADY CAI CTA AAA ATY TTR ATW CCD GT 3’ (modified from primer C1-J-2441 in Simon et al. 1994) using PCR conditions in Kaartinen et al. (2010). Gallwasp cytb sequences were amplified using primers CB1/CB2 (Jermiin and Crozier, 1994), using PCR conditions in Stone et al. (2009). For both parasitoids and gallwasps, the 28S D2 region was amplified using primers D2F/D2R (Heraty et al. 2004), using PCR conditions in Stone et al. (2009). PCRs were performed in 20µL reactions with the following final concentrations of each reagent: 1x PCR buffer; 2mM MgCl_2_ for cytb,1.5mM MgCl_2_ for COI and D2; 0.2µM of each primer; 125µM of each dNTP for COI, 200µM of each dNTP for cytb, and 250µM of each dNTP for D2; 1mg/mL BSA and 0.5U/µL polymerase (Bioline). Post-PCR cleanup used a SAP/ExoI protocol, and fragments were sequenced in both directions using BigDye v3.1 terminator chemistry on an ABI 3731XL capillary sequencer (Life Technologies/Thermofisher). Base calls were confirmed by eye in Sequencher 5.4.6 (Gene Codes Corporation).

Sequences were aligned using MAFFT online (Katoh et al. 2019) with G-INS-i specified as the iterative refinement method. Sequences for the coding COI and cytb mitochondrial genes were unambiguously aligned at 698 bp and 433 bp, respectively. However, because some previously published COI sequences were not amplified using the same COI_pf1/COI_2413d primer pair, CO1 sequences in the alignment ranged in length from 451 to 698 bp. D2 alignments were edited by removal of indels that appeared in a single specimen and of regions that were difficult to align unambiguously, giving final alignment lengths of 566 bp for gallwasps and 606 bp for parasitoids. Genbank accession numbers for all sequences are provided in the Supplemental PDF, table S2.

#### Selection of partition substitution models

Initial substitution models were estimated by analysis of a concatenated data matrix totalling 999 bp for gallwasps and 1304bp for parasitoids using the -m TEST command (Kalyaanamoorthy et al. 2017) in IQ-TREE v2.1.2 (Minh et al. 2020). The concatenated data matrix was partitioned by codon position for COI and cytb, with a separate partition for 28S D2. The best BIC models of respective partitions are listed in table A2.

We used the StarBEAST2 module (Ogilvie et al., 2017) implemented in BEAST2 (Bouckaert et al., 2019) to generate a species tree for each trophic level. Because the substitution models available in StarBEAST2 are limited to JC69, HKY, TN93 and GTR, we adopted the following procedure to select the closest suitable option to the IQ-TREE-inferred model (table A2). First, we reduced the complexity of the rate matrix for specific types of transition and transversion in each partition. We examined the empirical frequencies of transition and transversion types present using the pairwise base frequencies command in PAUP* 4.0 (Swofford 2003), and where specific transition or transversion types had very low frequencies, we assigned them the same rates as other types with higher frequencies, following recommendations in the MrBayes manual v3.2. Second, since the base frequencies parameter F is not an option in StarBEAST2, we excluded it from simplified models (table A2). Third, because partitions for gallwasp cytb codon positions 1 and 2 and parasitoid 28S D2 show a proportion of conserved sites (table A3), we incorporated proportion of invariant sites (I) as an additional parameter in the substitution models for these partitions. These substitution models were used in initial runs in StarBEAST2 with the following specifications: mitochondrial gene ploidy set to 0.5, population model: constant population size, phylogenetic model: birth-death, chain length set to 100 million generations and sampled every 12,500 generations. Because MCMCglmm requires ultrametric phylogenies (see below) we used a strict clock model.

Initial analysis of the 28S D2 gene tree of oak gallwasps resulted in multiple unresolved polytomies, with knock-on effects on the resolution of the two-gene species tree. Because the D2 gene alignment of oak gallwasps has 87.9% identical sites (table A3), we judged that inclusion only of the parameter I for rate heterogeneity across sites might not adequately account for between-site variability. We therefore respecified the substitution model for this gene to include gamma-distributed rate variation (G), with 4 gamma categories. This model yielded a better-resolved bifurcating tree, and the G parameter was therefore retained for downstream analyses. The final substitution models and rate parameters for all genes and partitions are shown in table A2.

#### Imposition of phylogenetic constraints and clock calibration

Initial analysis of the parasitoid COI data found the family Eulophidae to be paraphyletic, while the D2 gene tree for the same taxa supports monophyly. This discordance of gene tree topologies results in Eulophidae being paraphyletic in the species tree. Because more in-depth phylogenetic analyses support monophyly of Eulophidae (e.g. Burks et al. 2011; Munro et al. 2011; Rasplus et al. 2020), we constrained Eulophidae to be monophyletic in downstream analyses (see descriptions of monophyly settings below). All other parasitoid families in our dataset were recovered as monophyletic.

We calibrated molecular clocks for the gallwasp and parasitoid species trees to identify the timescales involved for phylogenetic patterns inferred in MCMCglmm. Node ages were calibrated in StarBEAST2 using estimates of node ages and 95% confidence intervals (CI) inferred in previous more extensive analyses by Peters et al. (2018) for parasitoids and by Blaimer et al. (2020) for gallwasps, following guidance for most recent common ancestor (MRCA) prior specification and setting of monophyletic group constraints in the Taming the BEAST online resource (https://taming-the-beast.org/tutorials/Introduction-to-BEAST2/; Barrido-Sottani et al. 2018). To allow date calibration using shared nodes in Peters et al. (2018) for the parasitoid species tree, we selected an MRCA prior and imposed monophyly for Eulophidae, Eupelmidae, Eurytomidae and Torymidae. To allow date calibration using shared nodes in Blaimer et al. (2020) for the oak gallwasp species tree, the same MRCA prior and monophyletic group constraint approach was applied to species in the lucidus clade (*Andricus grossulariae*, *A. lucidus*, *A. seckendorffi*) as defined in Stone et al. (2009), and to all species of oak gallwasps. A lognormal distribution (Ho and Phillips 2009) was applied for node age inference. The specific shared node ages in Peters et al. (2018) used in our clock calibration were node 18 (86 mya; CI: 62–146 mya) for the MRCA of parasitoids in our dataset, node 25 (40 mya; CI: 21–71 mya) for Torymidae, node 28 (48 mya; CI: 25–86 mya) for Eupelmidae, and node 36 (23 mya; CI: 11–40 mya) for Eurytomidae. The specific shared node ages in Blaimer et al. (2020) for gallwasps were node 129 (75.6 mya; CI: 50.5–106.9 mya) for Cynipini, and node 150 (17.2 mya; CI: 6.3–30.1 mya) for the lucidus clade. In StarBEAST2 monophyletic group priors, the minimum age CI (see above) was entered as the offset value for each monophyletic group, M was entered as the mean difference between the maximum CI and the minimum CI, and S was tuned such that the 2.5%–97.5% interquantile range matched the estimated CI values in Peters et al. (2018) and Blaimer et al. (2020).

#### Selection of appropriate population and clock models

Our MCMCglmm analyses require an ultrametric tree of each trophic level, which requires a strict clock model. To assess the suitability of such an assumption we compared the strict clock model to alternatives for the parasitoid and gallwasp sequence data. We used nested sampling (Skilling, 2006; Maturana et al., 2018) to estimate relative support for the best combination of the two population models (Analytical Population Size Integration *versus* Constant populations) and clock models (Strict Clock *versus* Uncorrelated Lognormal) available in BEAST2, following the online instructions in Taming the BEAST (Barido-Sottani et al., 2018; https://taming-the-beast.org/tutorials/NS-tutorial/). We first set up the substitution models for each partition as well as the values for time calibration, as described above, and ran models with the four possible population and clock model combinations, with chainLength=500,000, particleCount=1 and subChainLength=5,000. The difference in log of marginal likelihood between models was used to calculate Bayes Factors (BF) following Kass and Raftery (1995). We first compared the log of marginal likelihood between models to find the best two models, and then further compared the best two models using BF. Where the BF < 2*√(SD1^2^+SD2^2^) between the best two models, we increased the particleCount for a longer sampling, following the aforementioned online instructions for nested sampling in Taming the BEAST (Barido-Sottani et al., 2018).

For the gallwasp data, the alternative population models combined with the Uncorrelated Lognormal clock model are indistinguishable from each other, but both have higher log marginal likelihood than either of the two population models combined with the Strict Clock model. In parasitoids, the two clock models combined with Analytical Population Size Integration population model could not be differentiated from each other but were better than the two clock models combined with Constant populations. We carried out additional longer analyses to further resolve log marginal likelihood estimates for the two better supported model combinations for each dataset (two population models with Uncorrelated Lognormal clock model for gallwasps, and the combinations of two clock models with Analytical Population Size Integration population model for parasitoids). We used the formula SD = sqrt(H/N) where SD is the standard deviation of the log marginal likelihood estimate for a model, H was taken from the information value in the first run of the model combination with the best log of marginal likelihood, and N is the number of particles (Barido-Sottani et al., 2018). The target SD values using this approach were 6 and 5 for gallwasps and parasitoids respectively, which made N = 23 in both trophic levels for further model selection processes. chainLength and subChainLength were as for the initial runs. Model comparison using the procedure above then favored Analytical Population Size Integration + Uncorrelated Lognormal for gallwasps (table A4), and Analytical Population Size Integration + Strict Clock for parasitoids (table A5).

In the final phylogenetic analyses we applied all the estimated models and divergence times for species tree inference in StarBEAST2. Analyses were run for 100 million generations and sampled every 12,500 generations for both parasitoids and gallwasps. Data convergence was examined in Tracer 1.7.2 (Rambaut et al., 2018) using the log file. The maximum clade credibility trees used in the MCMCglmm analyses were generated from the last 20% of generations using common ancestor height in TreeAnnotator (included in BEAST 2.6.6), and the tree was visualised in FigTree 1.4.4 (https://github.com/rambaut/figtree/releases).

**Table A2.**
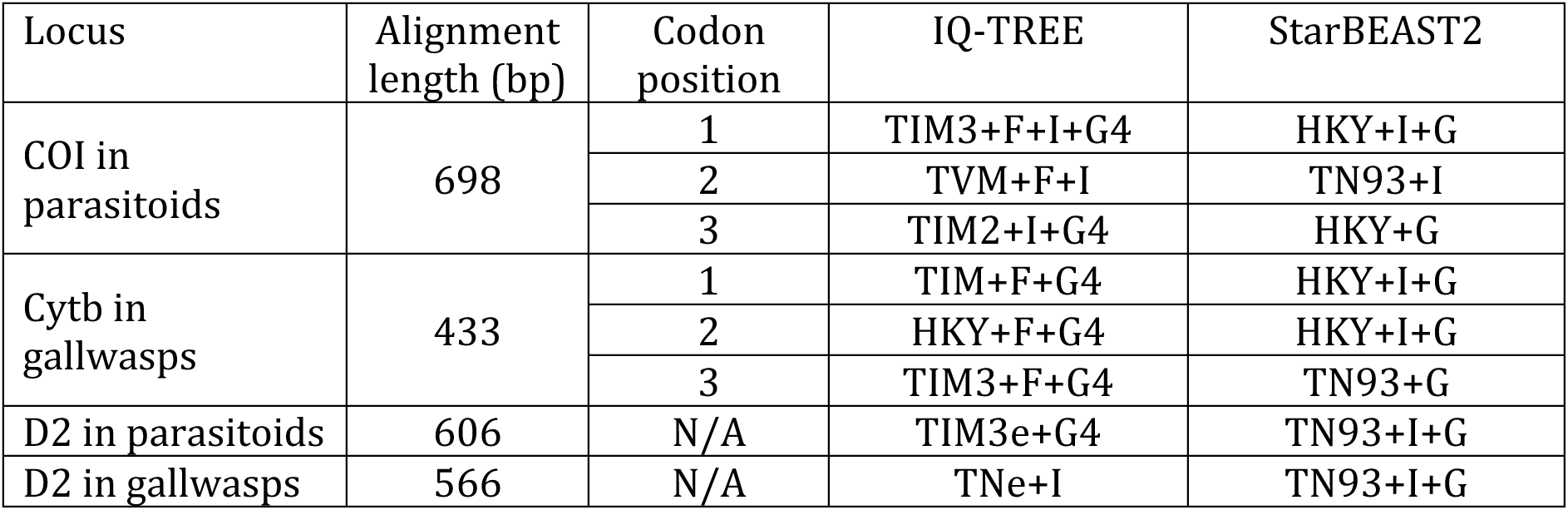
Substitution models for each gene and codon position estimated in IQ-TREE, and the simplified nearest equivalents used in StarBEAST2 analyses. N/A for partition indicates that a gene fragment was treated as a single partition.

| Locus | Alignment length (bp) | Codon position | IQ-TREE | StarBEAST2 |
| --- | --- | --- | --- | --- |
| COI in parasitoids | 698 | 1 | TIM3+F+I+G4 | HKY+I+G |
|  |  | 2 | TVM+F+I | TN93+I |
|  |  | 3 | TIM2+I+G4 | HKY+G |
| Cytb in gallwasps | 433 | 1 | TIM+F+G4 | HKY+I+G |
|  |  | 2 | HKY+F+G4 | HKY+I+G |
|  |  | 3 | TIM3+F+G4 | TN93+G |
| D2 in parasitoids | 606 | N/A | TIM3e+G4 | TN93+I+G |
| D2 in gallwasps | 566 | N/A | TNe+I | TN93+I+G |

**Table A3.** Partition sizes by genes and codon positions, and the number of conserved and variable sites within each partition. Position

| Locus | Codon position | Length (bp) | Conserved | Variable | % Invariant sites |
| --- | --- | --- | --- | --- | --- |
| COI in parasitoids | 1 | 232 | 154 | 78 | 66.3 |
|  | 2 | 233 | 199 | 34 | 85.4 |
|  | 3 | 233 | 11 | 222 | 4.7 |
| Cytb in gallwasps | 1 | 144 | 98 | 46 | 68.0 |
|  | 2 | 144 | 121 | 23 | 84.0 |
|  | 3 | 145 | 18 | 127 | 12.4 |
| D2 in parasitoids | N/A | 606 | 426 | 180 | 70.2 |
| D2 in gallwasps | N/A | 566 | 498 | 68 | 87.9 |

**Table A4.** Model log marginal likelihoods used in Bayes Factor (BF)-based model selection for the gallwasp sequence dataset. In each run, the models are ranked by log marginal likelihood. Support for the top two ranked models is compared using BF, with the assessment of meaningful difference based on the BF ≥ 2*(SD1+SD2) where SD1 and SD2 are the standard deviations of the log marginal likelihoods for the 2 models being compared. N is the number of particles and H is the information content.

| | Model | N | H | SD | log marginal likelihood | BF (ML1–ML2) | $2*\sqrt{(SD1^2+SD2^2)}$ | Model selection result |
| --- | --- | --- | --- | --- | --- | --- | --- | --- |
| First run | Analytical Population Size Integration + Uncorrelated clock lognormal | 1 | 818.9 | 28.6 (SD1) | -7545.1 | 38.6 | 83.62 | BF < $2*\sqrt{(SD1^2+SD2^2)}$<br>Models are indistinguishable |
|  | Constant populations + Uncorrelated clock lognormal | 1 | 928.3 | 30.5 (SD2) | -7583.7 |  |  |  |
|  | Analytical Population Size Integration + Strict clock | 1 | 834.3 | 28.9 (SD3) | -7789.5 |  |  |  |
|  | Constant populations + Strict clock | 1 | 874.8 | 29.6 (SD4) | -7683.1 |  |  |  |
| Second run | Analytical Population Size Integration + Uncorrelated clock lognormal | 23 | 643.5 | 5.3 (SD1) | -7307.5 | 167.9 | 15.86 | BF > $2*\sqrt{(SD1^2+SD2^2)}$<br>Analytical Population Size Integration + Uncorrelated lognormal clock is overwhelming supported |
|  | Constant populations + Uncorrelated clock lognormal | 23 | 812.0 | 5.9 (SD2) | -7475.3 |  |  |  |

**Table A5.** Model log marginal likelihoods used in Bayes Factor (BF)-based model selection for the parasitoid sequence dataset. In each run, the models are ranked by log marginal likelihood. Support for the top two ranked models is compared using BF, with the assessment of meaningful difference based on the BF ≥ 2*(SD1+SD2) where SD1 and SD2 are the standard deviations of the log marginal likelihoods for the 2 models being compared. N is the number of particles and H is the information content.

| | Model | N | H | SD | log marginal likelihood | BF (ML1–ML2) | $2 \cdot \sqrt{(SD1^2 + SD2^2)}$ | Model selection result |
| --- | --- | --- | --- | --- | --- | --- | --- | --- |
| First run | Analytical Population Size Integration + Strict clock | 1 | 556.2 | 23.6 (SD1) | -15134.1 (ML1) | 11.0 | 65.63 | $BF < 2 \cdot \sqrt{(SD1^2 + SD2^2)}$<br>Models are indistinguishable |
|  | Analytical Population Size Integration + Uncorrelated clock lognormal | 1 | 519.4 | 22.8 (SD2) | -15145.1 (ML2) |  |  |  |
|  | Constant populations + Strict clock | 1 | 456.3 | 21.4 (SD3) | -15352.28 (ML3) |  |  |  |
|  | Constant populations + Uncorrelated clock lognormal | 1 | 517.7 | 30.5 (SD4) | -15311.66 (ML4) |  |  |  |
| Second run | Analytical Population Size Integration + Strict clock | 23 | 441.6 | 4.4 (SD1) | -15002.1 (ML1) | 41.5 | 12.44 | $BF > 2 \cdot \sqrt{(SD1^2 + SD2^2)}$<br>Analytical Population Size Integration+Strict clock is overwhelmingly supported |
|  | Analytical Population Size Integration + Uncorrelated clock lognormal | 23 | 455.3 | 4.4 (SD2) | -15043.6 (ML2) |  |  |  |

## Supplementary Methods

**Figure S1.**
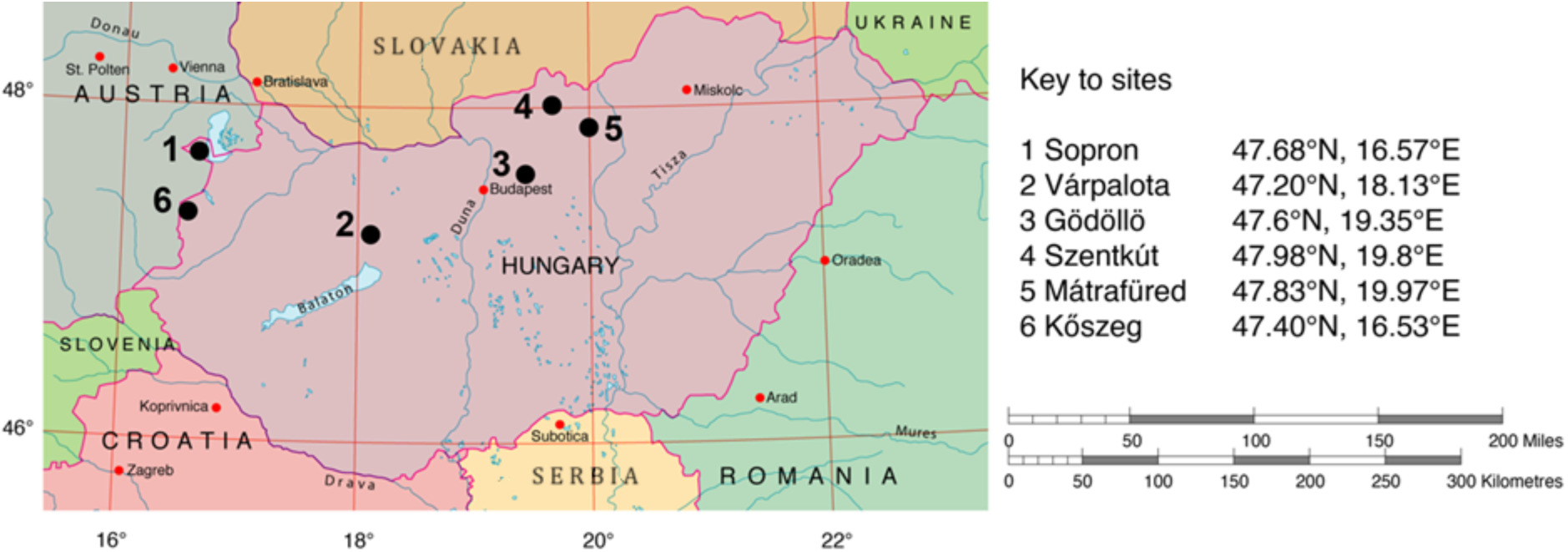
Sample site locations.

### Taxonomic resolution

The Linnaean (morphology-based) taxonomy of both trophic levels in our datasets is well established (Richard R. Askew et al. 2013; Roskam 2019). However, since our datasets were generated, cryptic taxa have been demonstrated in four of the parasitoid species: Bootanomyia (Megastigmus) dorsalis and B. synophri (Nicholls et al. 2018; 2010), and Torymus cyaneus and T. flavipes (Kaartinen et al. 2010; Gil-Tapetado et al. 2022). All four Linnaean species attack multiple host galls, and B. dorsalis and T. flavipes are two of the more generalist parasitoids in Western Palaearctic oak cynipid galls (Richard R. Askew et al. 2013)). The impact of lumping cryptic species on our inference depends critically on how links recorded for the Linnaean species are divided among them. If cryptic sister parasitoid species attack similar sets of related host galls, we would expect increased support for the Cophylogenetic interaction. DNA barcode-based analyses have shown cryptic species within Bootanomyia (Megastigmus) dorsalis and T. flavipes to still be generalists (Kaartinen et al. 2010; Nicholls et al. 2018; 2010). In the absence of directional predictions for potential errors associated with cryptic taxa, we interpret our results at face value while noting that consideration of cryptic taxa is an issue in all trophic network analyses (Pringle and Hutchinson 2020).

**Table S1.** Sampled gall-types in Spring (S) and Autumn (A) datasets, with details of the number of galls reared, the number of parasitoid individuals that emerged from the reared galls, and the number of different parasitoid links.

| Cynipid species and generation | # galls | # parasitoids | # links |
| --- | --- | --- | --- |
| Sexual generation dataset |  |  |  |
| *Andricus conificus* (S) (Hartig, 1843) | 340 | 113 | 18 |
| *Andricus curvator* (S) Hartig, 1840 | 644 | 146 | 12 |
| *Andricus grossulariae* (S) Giraud, 1859 | 955 | 224 | 16 |
| *Andricus inflator* (S) (Hartig, 1840) | 51 | 2 | 2 |
| *Andricus lucidus* (S) (Hartig, 1843) | 32 | 85 | 4 |
| *Andricus multiplicatus* (S) Giraud, 1859 | 418 | 315 | 14 |
| *Andricus quadrilineatus* (A) (Hartig, 1840) | 368 | 6 | 4 |
| *Andricus quercuscalicis* (S) (Burgsdorf 1783) | 41 | 3 | 1 |
| *Andricus quercusramuli* (S) (Linnaeus, 1761) | 248 | 1940 | 14 |
| *Andricus schroeckingeri* (S) Wachtl, 1876 | 600 | 64 | 18 |
| *Andricus seckendorffi* (S) Wachtl (=A. *vindobonensis* Müllner, 1901) | 721 | 27 | 7 |
| *Andricus singularis* (S) Mayr, 1870 | 299 | 95 | 14 |
| *Andricus superfetationis* (S) (Giraud, 1859) (=A. *crispator* Tschek, 1871) | 122 | 261 | 12 |
| *Biorhiza pallida* (S) (Olivier, 1791) | 1185 | 12508 | 29 |
| *Chilaspis nitida* (S) (Giraud, 1882) | 330 | 1297 | 23 |
| *Cynips disticha* (S) Hartig, 1840 | 67 | 33 | 5 |
| *Cynips divisa* (S) Hartig, 1840 | 9 | 6 | 3 |
| *Cynips longiventris* (S) Hartig, 1840 | 3 | 2 | 1 |
| *Cynips quercusfolii* (S) (Linnaeus, 1758) | 13 | 3 | 2 |
| *Dryocosmus cerriphilus* (S) (Giraud, 1859) | 2 | 0 | 0 |
| *Neuroterus albipes* (S) (Schenck, 1863) | 6 | 0 | 0 |
| *Neuroterus anthracinus* (S) (Curtis, 1838) | 565 | 195 | 18 |
| *Neuroterus numismalis* (S) Geoffroy in Fourcroy, 1785 | 777 | 121 | 11 |
| *Neuroterus politus* (S) Hartig, 1840 | 8 | 0 | 0 |
| *Neuroterus quercusbaccarum* (S) (Linnaeus, 1758) | 846 | 129 | 5 |
| *Pseudoneuroterus saliens* (S) (Kollar, 1857) | 3141 | 635 | 22 |
| Asexual generation dataset |  |  |  |
| *Andricus amblycerus* (A) (Giraud, 1859) | 271 | 18 | 6 |
| *Andricus aries* (A) (Giraud, 1859) | 54 | 2 | 2 |
| *Andricus caliciformis* (A) (Giraud, 1859) | 151 | 45 | 7 |
| *Andricus caputmedusae* (A) (Hartig, 1843) | 592 | 478 | 20 |
| *Andricus conglomeratus* (A) (Giraud, 1859) | 624 | 124 | 12 |
| *Andricus conificus* (A) (Hartig, 1843) | 293 | 52 | 7 |
| *Andricus coriarius* (A) (Hartig, 1843) | 775 | 2332 | 17 |
| *Andricus coronatus* (A) (Giraud, 1859) | 189 | 129 | 9 |
| *Andricus corruptrix* (A) (Schlechtendal, 1870) | 72 | 8 | 5 |
| *Andricus dentimitratus* (A) (Rejtő, 1887) | 3 | 0 | 0 |
| *Andricus foecundatrix* (A) (Hartig, 1840) | 243 | 9 | 5 |
| *Andricus galeatus* (A) (Giraud, 1859) | 260 | 32 | 8 |
| *Andricus gemmeus* (A) (Giraud, 1859) | 62 | 7 | 3 |
| *Andricus glandulae* (A) (Hartig, 1840) | 11 | 2 | 2 |
| *Andricus glutinosus* (A) (Giraud, 1859) | 576 | 108 | 11 |
| *Andricus grossulariae* (A) Giraud, 1859 | 435 | 1292 | 20 |
| *Andricus hartigi* (A) (Hartig, 1843) | 116 | 36 | 7 |
| *Andricus hungaricus* (A) Hartig, 1843 | 462 | 607 | 13 |
| *Andricus hystrix* (A) Kieffer, 1897 | 166 | 8 | 3 |
| *Andricus infectorius* (A) (Hartig, 1843) | 1197 | 176 | 12 |
| *Andricus inflator* (A) (Hartig, 1840) | 57 | 4 | 2 |
| *Andricus kollari* (A) (Hartig, 1843) | 532 | 473 | 20 |
| *Andricus lignicolus* (A) (Hartig, 1840) | 928 | 251 | 12 |
| *Andricus lucidus* (A) (Hartig, 1843) | 647 | 662 | 15 |
| *Andricus mitratus* (A) (Mayr, 1870) | 3 | 1 | 1 |
| *Andricus paradoxus* (A) (Radoszkowski, 1866) | 31 | 0 | 0 |
| *Andricus polycerus* (A) (Giraud, 1859) | 219 | 17 | 4 |
| *Andricus quercuscalicis* (A) (Burgsdorf 1783) | 858 | 414 | 16 |
| *Andricus quercustozae* (A) (Bosc, 1792) | 769 | 54 | 7 |
| *Andricus seckendorffi* (A) Wachtl | 9 | 6 | 2 |
| *Andricus solitarius* (A) (Fonscolombe, 1832) | 75 | 16 | 4 |
| *Andricus superfetationis* (A) (Giraud, 1859) | 39 | 2 | 2 |
| *Andricus truncicolus* (A) (Giraud, 1859) | 1 | 0 | 0 |
| *Aphelonyx cerricola* (A) (Giraud, 1859) | 532 | 153 | 13 |
| *Callirhytis glandium* (A) (Giraud, 1859) | 697 | 61 | 8 |
| *Cerroneuroterus lanuginosus* (A) (Giraud, 1859) | 574 | 37 | 11 |
| *Chilaspis nitida* (A) (Giraud, 1882) | 959 | 88 | 11 |
| *Cynips cornifex* (A) Hartig, 1843 | 492 | 49 | 8 |
| *Cynips disticha* (A) Hartig, 1840 | 426 | 43 | 12 |
| *Cynips divisa* (A) Hartig, 1840 | 316 | 40 | 7 |
| *Cynips longiventris* (A) Hartig, 1840 | 201 | 61 | 12 |
| *Cynips quercus* (A) (Fourcroy, 1785) | 2800 | 438 | 17 |
| *Cynips quercusfolii* (A) (Linnaeus, 1758) | 1519 | 363 | 19 |
| *Dryocosmus cerriphilus* (A) (Giraud, 1859) | 8 | 0 | 0 |
| *Neuroterus albipes* (A) (Schenck, 1863) | 106 | 6 | 3 |
| *Neuroterus anthracinus* (A) (Curtis, 1838) | 1438 | 77 | 12 |
| *Neuroterus numismalis* (A) Geoffroy in Fourcroy, 1785 | 726 | 35 | 5 |
| *Neuroterus quercusbaccarum* (A) (Linnaeus, 1758) | 2001 | 97 | 11 |
| *Pseudoneuroterus macropterus* (A) (Hartig, 1843) | 195 | 181 | 13 |
| *Pseudoneuroterus saliens* (A) (Kollar, 1857) | 2247 | 117 | 14 |
| *Trigonaspis megaptera* (A) (Panzer, 1801) | 246 | 2 | 2 |
| *Trigonaspis synaspis* (A) (Hartig, 1841) | 644 | 22 | 7 |

**Table S2.**
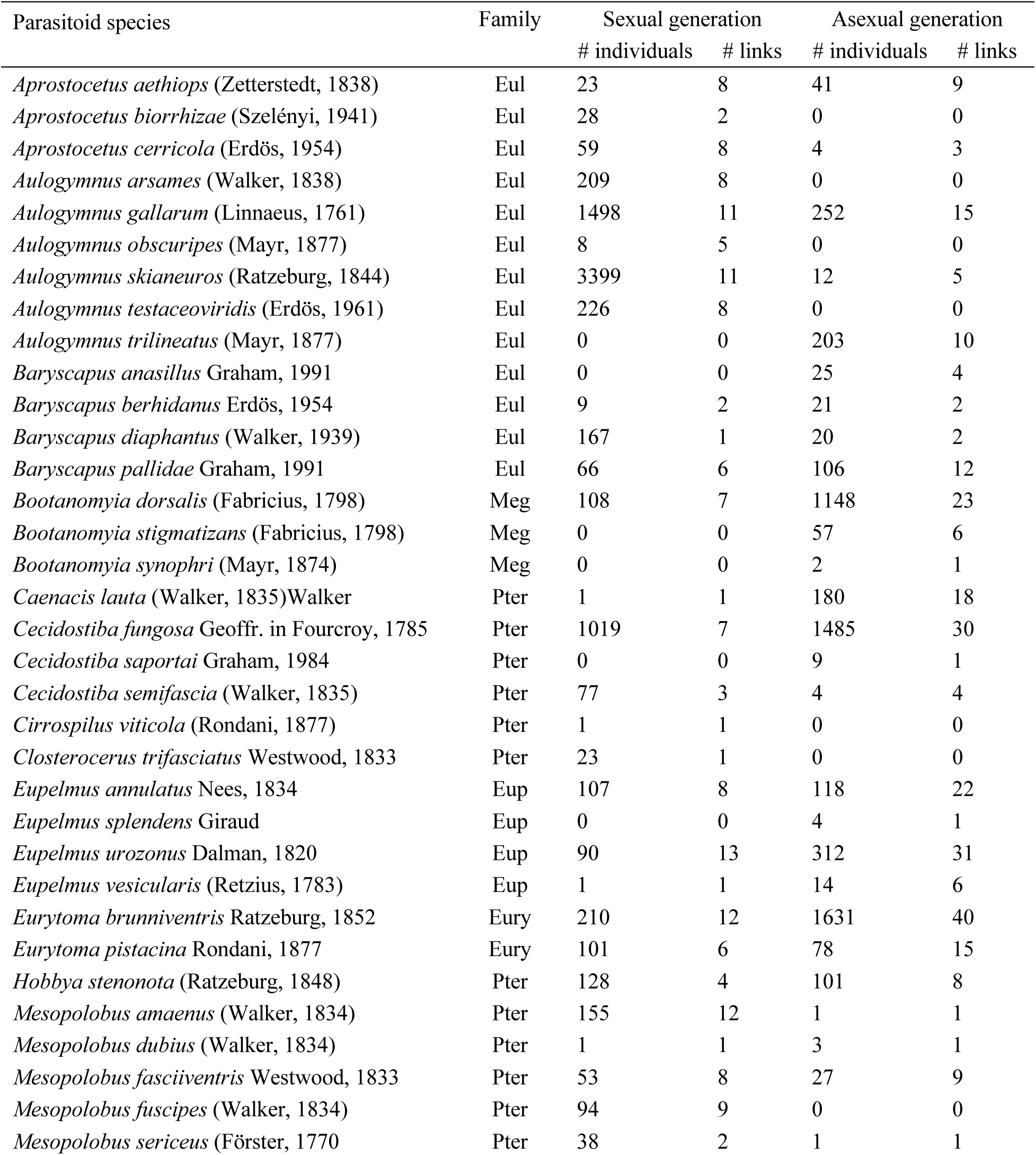
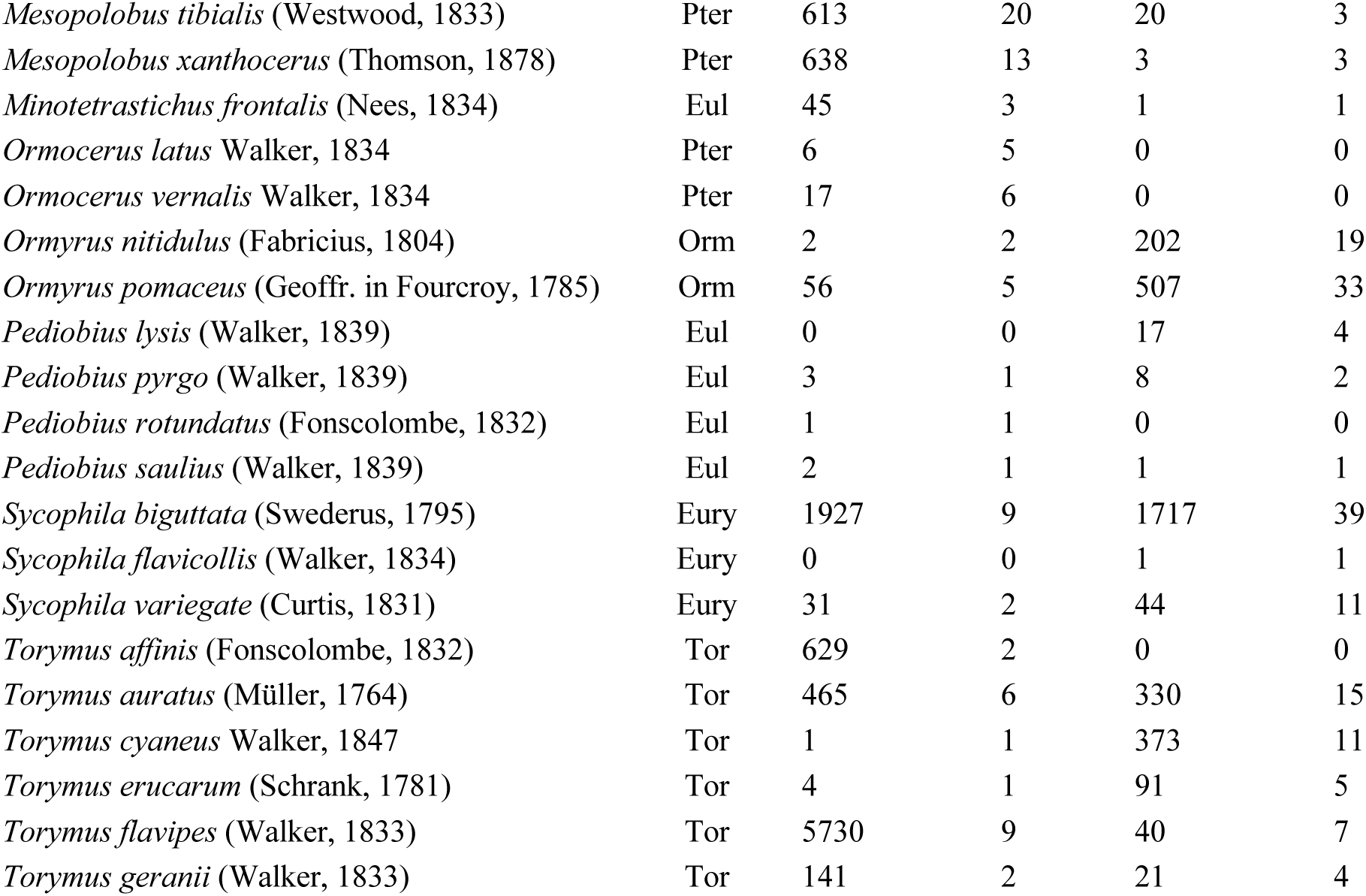
Alphabetic list of recorded parasitoid species, with details of the number of individuals and the number of different host gall-type links from each host gall generation. Parasitoid family names are abbreviated as follows: Eulophidae (Eul), Eupelmidae (Eup), Eurytomidae (Eury), Megastigmidae (Meg), Ormyridae (Orm), Pteromalidae (Pter), Torymidae (Tor). Species previously placed in the genus *Megastigmus* are here given their currently recognised names in the genus *Bootanomyia*. Since the datasets in our study were generated, molecular studies have shown that three of the morphologically defined parasitoid species in our data (*Megastigmus dorsalis, Torymus cyaneus and T. flavipes*) comprise sets of cryptic sister taxa (Kaartinen et al. 2010; Nicholls et al. 2010, 2018). Our analyses retain the original morphological classification as most of the original specimens are unavailable for sequence-based verification.

**Table S3.**
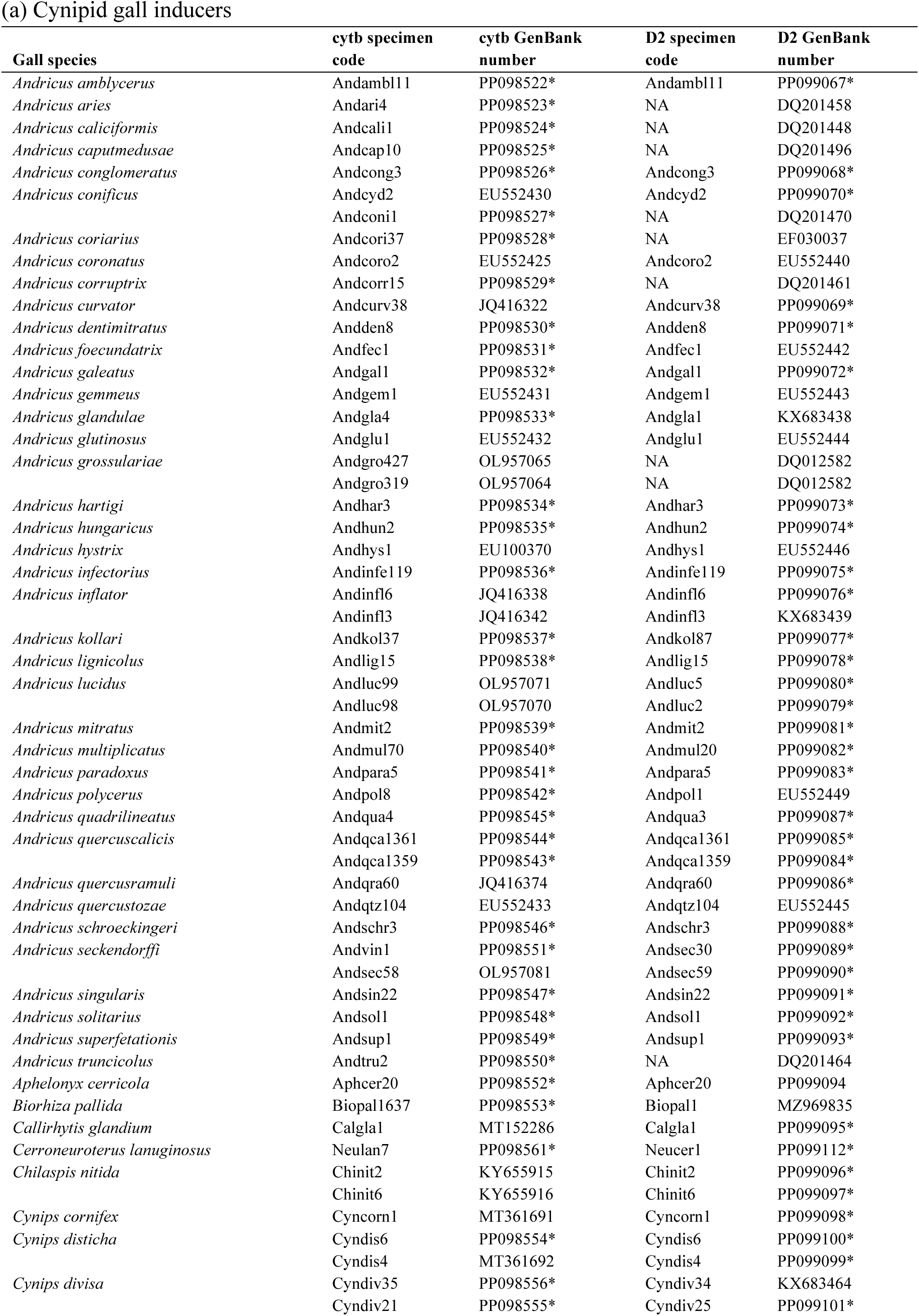
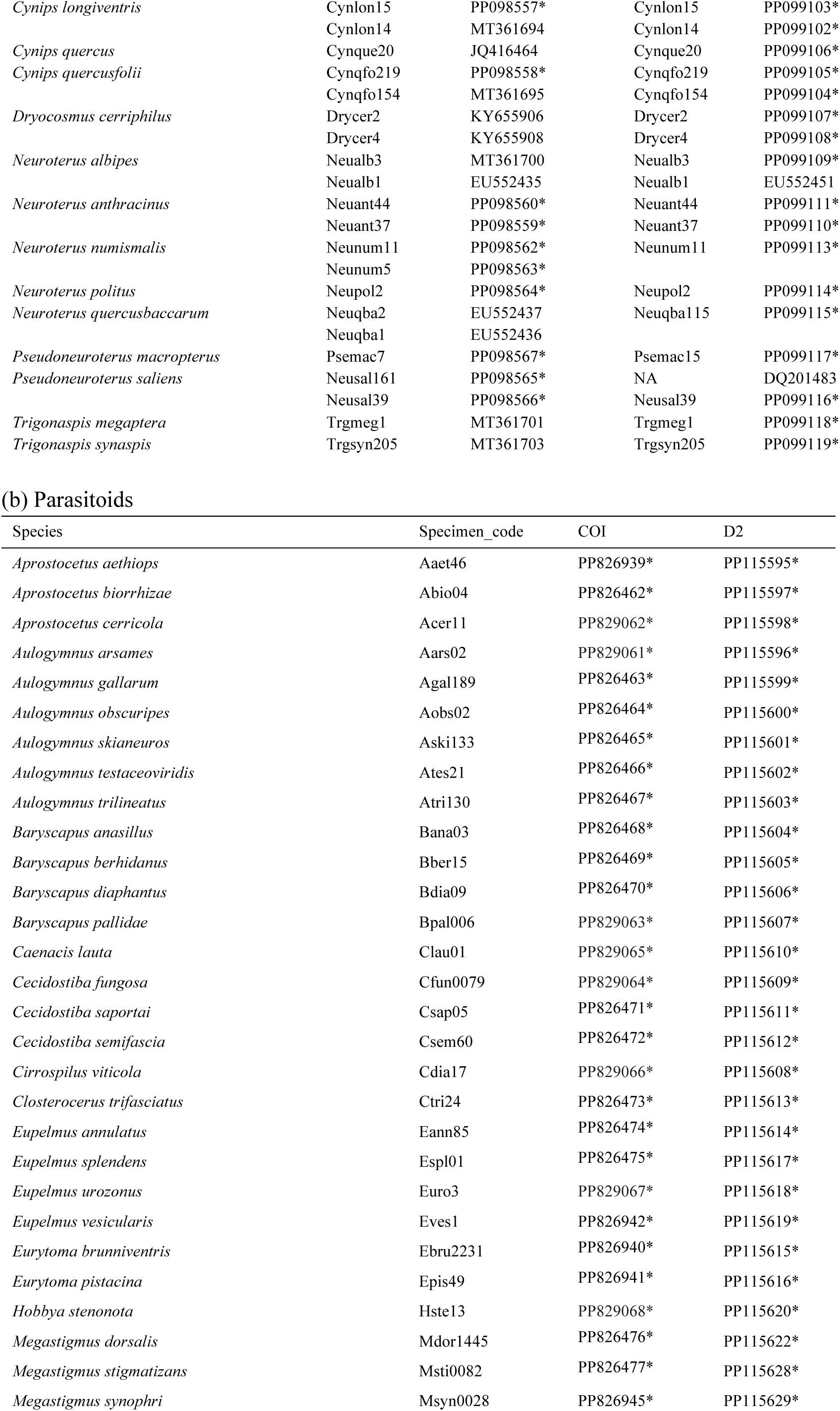
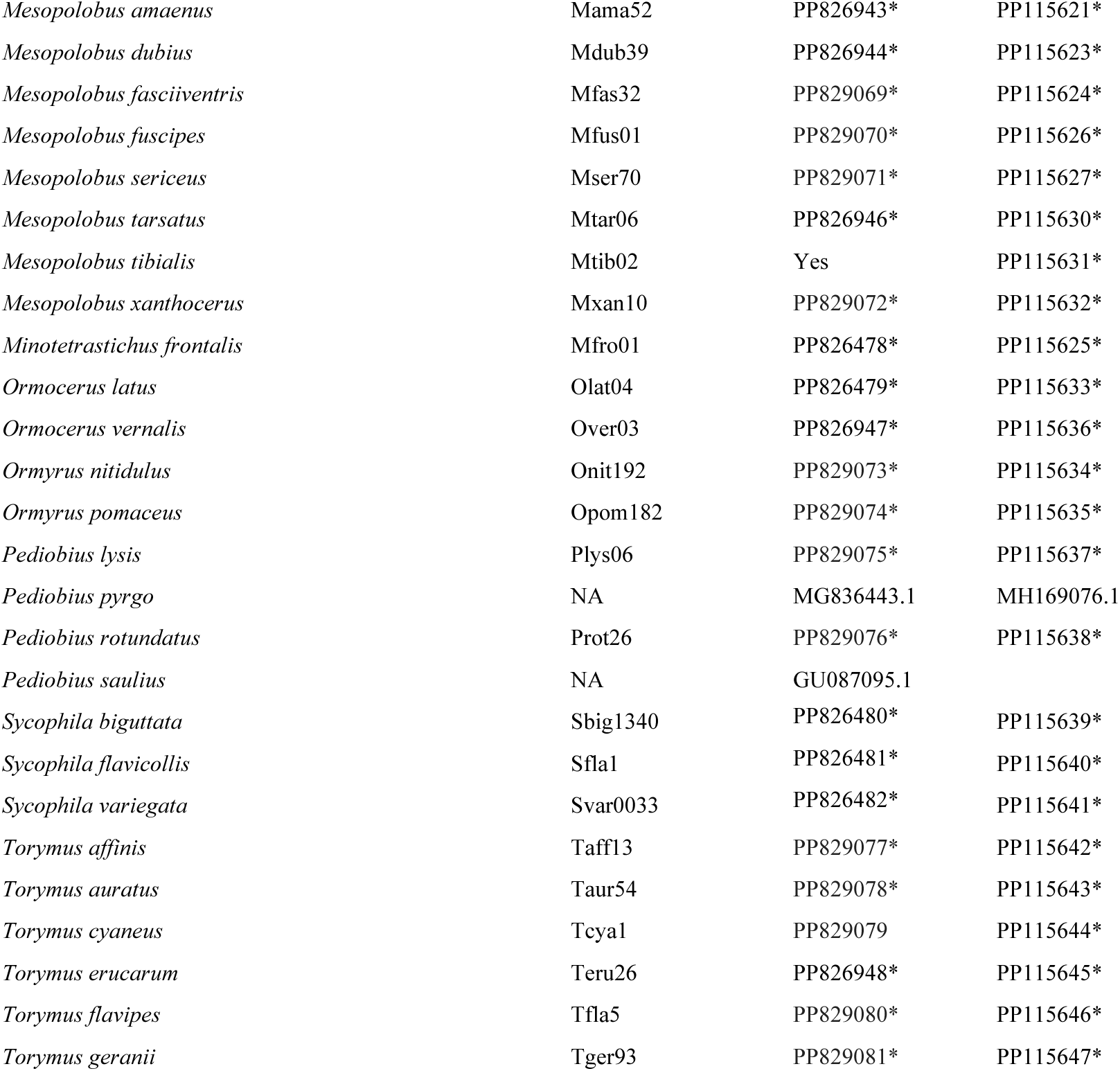
GenBank accession numbers and internal specimen codes (where applicable) for (a) cynipid gall inducers and (b) parasitoid species used in phylogeny construction. New sequences are indicated with an asterisk.

## Supplementary Results

**Tables S4 and S5** show intraclass correlations (ICCs) for terms in alternate versions of the 4 core models (incidence and frequency-based models of sexual and asexual generation gall datasets). Each table shows results for the full model and one or two alternate models in which specific model terms are excluded. ICC values are presented as median/mode (and 95% credible interval) over 2000 MCMC point estimates. NA indicates a model term that cannot be fitted in the specified model. For ease of comparison across models, significant terms (i.e. where the lower 95% credible interval >0.01) are highlighted in each model. ICC_GLMM_ is the ICC for all fitted model terms (i.e. excluding the residual), analogous to an R^2^ for the model. In the Full model (option 1), interactions between taxa missing from an individual site are omitted as structural zeros. In the Full model (option 2) and Pooled sites models, we assumed that the full set of parasitoid species recorded in a given generation across all six sites was available to interact at each site, and all links absent from a single site and generation were given a value of zero.

**Table S4.** Sexual generation models.

| Model term | Incidence models |  |  |  | Frequency models |  |  |
| --- | --- | --- | --- | --- | --- | --- | --- |
|  | Full model<br>(option 1) | Without sample<br>size | Full model<br>(option 2) | Pooled sites<br>(option 2) | Full model<br>(option 1) | Full model<br>(option 2) | Pooled sites<br>(option 2) |
| Site | .014 / .000<br>(.000-.096) | .028 / .016<br>(.000-.147) | .012 / .000<br>(.000-.090) | NA | .027 / .016<br>(.000-.134) | .092 / .063<br>(.008-.374) | NA |
| Site:host interaction | .030 / .031<br>(.006-.069) | .073 / .066<br>(.028-.143) | .054 / .046<br>(.016-.104) | NA | .064 / .055<br>(.025-.122) | .139 / .119<br>(.062-.243) | NA |
| Site:parasitoid interaction | .001 / .000<br>(.000-.006) | .001 / .000<br>(.000-.008) | .001 / .000<br>(.000-.010) | NA | .002 / .000<br>(.000-.009) | .002 / .000<br>(.000-.008) | NA |
| Host species | .031 / .001<br>(.000-.142) | .020 / .001<br>(.000-.141) | .046 / .001<br>(.000-.140) | .055 / .001<br>(.000-.163) | .024 / .001<br>(.000-.157) | .055 / .001<br>(.000-.231) | .073 / .002<br>(.000-.270) |
| Parasitoid species | .007 / .000<br>(.000-.049) | .029 / .001<br>(.000-.096) | .023 / .000<br>(.000-.096) | .030 / .001<br>(.000-.135) | .040 / .039<br>(.000-.099) | .054 / .047<br>(.008-.116) | .112 / .084<br>(.026-.227) |
| Host phylogeny | .179 / .002<br>(.000-.471) | .316 / .258<br>(.000-.595) | .054 / .001<br>(.000-.351) | .086 / .002<br>(.000-.424) | .314 / .340<br>(.000-.579) | .288 / .003<br>(.000-.621) | .380 / .003<br>(.000-.688) |
| Parasitoid phylogeny | .019 / .000<br>(.000-.106) | .014 / .000<br>(.000-.098) | .043 / .001<br>(.000-.162) | .053 / .001<br>(.000-.212) | .015 / .001<br>(.000-.099) | .011 / .000<br>(.000-.081) | .027 / .001<br>(.000-.158) |
| Parasitoid:host species interaction | .207 / .224<br>(.095-.329) | .132 / .138<br>(.056-.227) | .209 / .211<br>(.103-.320) | NA | .143 / .159<br>(.068-.230) | .057 / .064<br>(.021-.100) | NA |
| Host phylogenetic interaction | .013 / .000<br>(.000-.077) | .022 / .001<br>(.000-.104) | .024 / .001<br>(.000-.115) | .084 / .002<br>(.000-.255) | .009 / .000<br>(.000-.070) | .005 / .000<br>(.000-.038) | .011 / .001<br>(.000-.077) |
| Parasitoid phylogenetic interaction | .133 / .123<br>(.000-.268) | .072 / .046<br>(.000-.173) | .143 / .125<br>(.000-.261) | .185 / .215<br>(.000-.358) | .047 / .000<br>(.000-.110) | .027 / .024<br>(.000-.061) | .053 / .040<br>(.000-.122) |
| Cophylogenetic interaction | .089 / .002<br>(.000-.281) | .065 / .002<br>(.000-.282) | .052 / .001<br>(.000-.251) | .054 / .002<br>(.000-.332) | .038 / .001<br>(.000-.192) | .012 / .000<br>(.000-.086) | .021 / .001<br>(.000-.177) |
| Residual | .171 / .161<br>(.097-.248) | .114 / .105<br>(.060-.178) | .201 / .207<br>(.121-.275) | .292 / .294<br>(.128-.457) | .185 / .237<br>(.052-.299) | .159 / .164<br>(.032-.254) | .250 / .257<br>(.092-.407) |
| ICC denominator | 25.157 / 22.228<br>(16.192-40.403) | 37.743 / 32.890<br>(22.131-68.044) | 21.299 / 20.794<br>(14.473-32.140) | 14.694 / 12.945<br>(8.108-27.657) | 30.150 / 27.075<br>(18.451-49.273) | 74.146 / 63.437<br>(40.433-143.656) | 34.182 / 30.792<br>(18.500-61.745) |
| ICC<sub>GLMM</sub> | .829 / .839 (.752-.903) | .886 / .895<br>(.822-.940) | .799 / .793<br>(.725-.879) | .708 / .706<br>(.543-.872) | .815 / .763<br>(.701-.948) | .841 / .836<br>(.746-.968) | .750 / .743<br>(.593-.908) |
| ICC<sub>PHY</sub> | .263 / .291<br>(.098-.443) | .194 / .173<br>(.060-.368) | .253 / .221<br>(.119-.435) | .383 / .409<br>(.157-.631) | .118 / .076<br>(.033-.261) | .057 / .050<br>(.014-.130) | .113 / .094<br>(.028-.253) |
| ICC<sub>REL-PHY</sub> | .561 / .581<br>(.311-.763) | .597 / .609<br>(.345-.834) | .550 / .563<br>(.302-.752) | NA | .460 / .445<br>(.186-.699) | .503 / .486<br>(.269-.768) | NA |

**Table S5.**
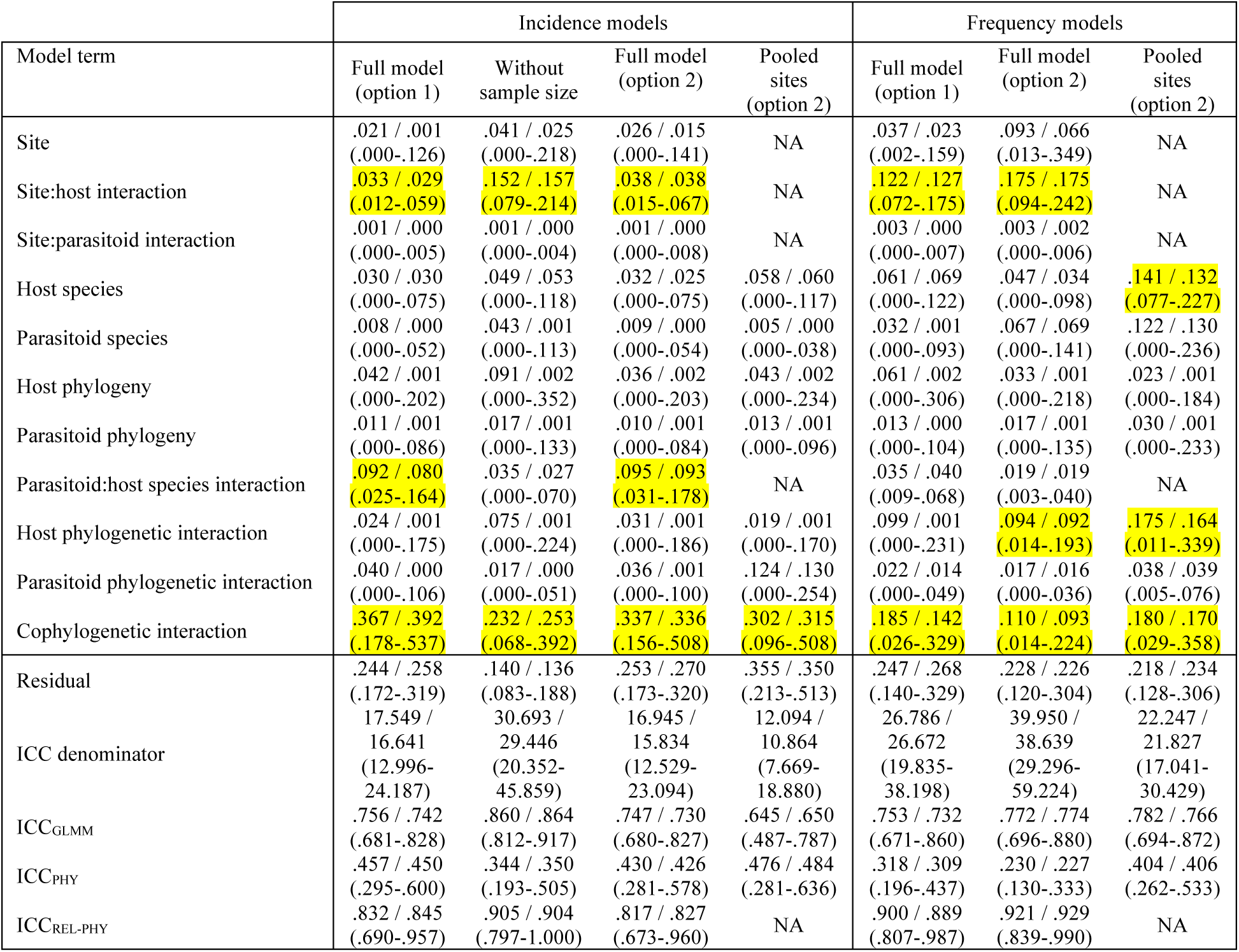
Asexual generation models

### Sensitivity of model results to reduced sampling intensity

**Figure S2.**
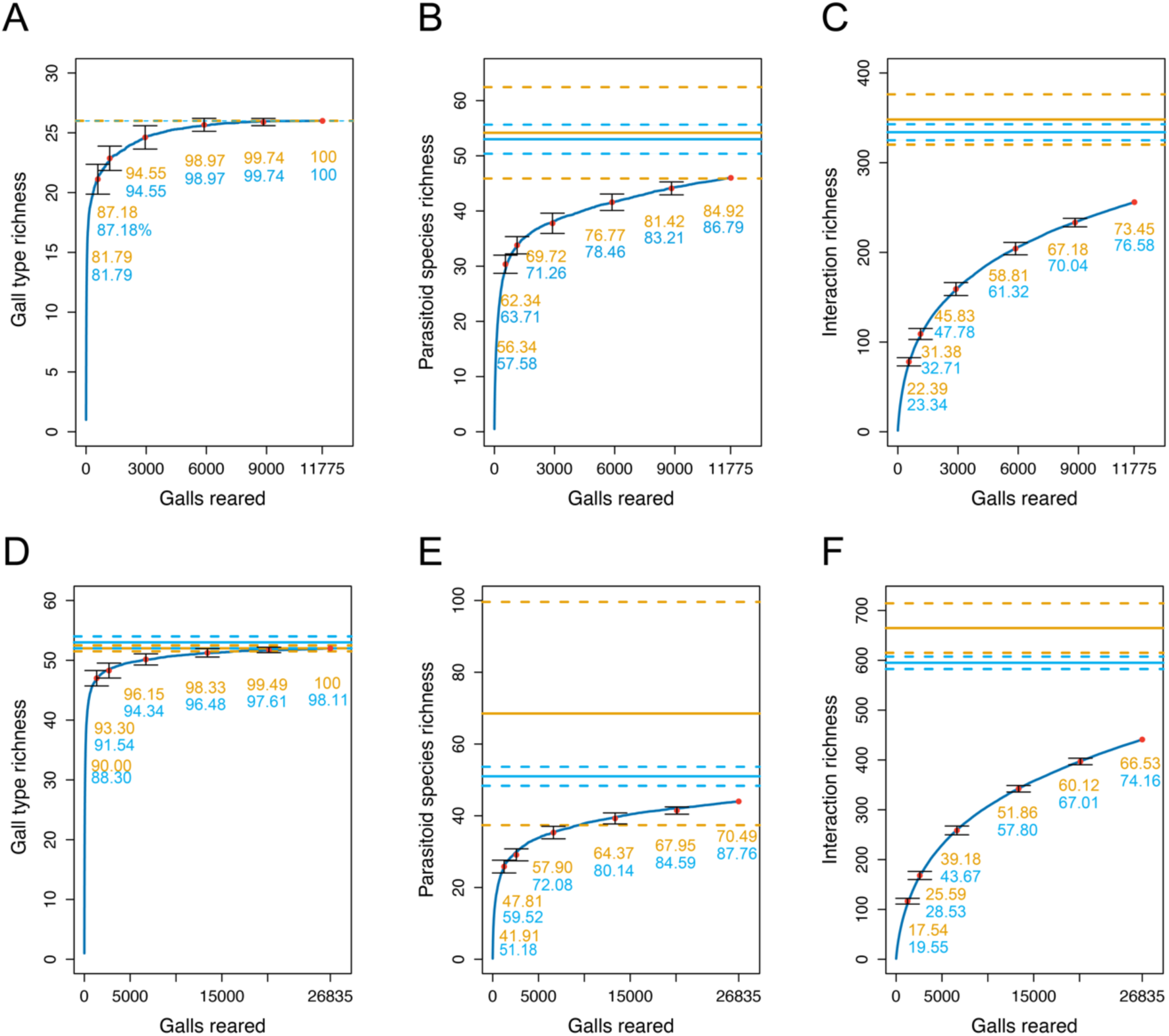
Accumulation curves in the sexual generation (A-C, top row) and asexual generation (D-F, bottom row) datasets for gall type richness (A, D), parasitoid species richness (B, E), and interaction richness (C, F). Horizontal lines indicate the Chao-2 (orange) and first order jackknife (blue) estimates of richness (with +/- 1 standard deviation) from the full datasets. From left to right the red points indicate the mean observed richness for 5%, 10%, 25%, 50%, and 75% subsets of the full data with bars for +/- 1 standard deviation, and the observed value for the full dataset. Values below points show their percentage relative to the Chao-2 (orange) and first order jackknife (blue) richness estimates.

